# The silent impact: codon usage bias and protein evolution in bacteria

**DOI:** 10.1101/2022.07.07.499116

**Authors:** Ana Filipa Moutinho, Adam Eyre-Walker

## Abstract

Bias in synonymous codon usage has been reported across all kingdoms of life. Evidence across species suggests that codon usage bias is often driven by selective pressures, typically for translational efficiency. These selective pressures have been shown to depress the rate at which synonymous sites evolve. We hypothesise that selection on synonymous codon use could also slow the rate of protein evolution if two amino acids have different preferred codons. We test this hypothesis by looking at patterns of protein evolution using polymorphism and substitution data in bacteria. We found that non-synonymous mutations that change from unpreferred to preferred codons are more common than the opposite, but only amongst codons that vary substantially in their preference level. Overall, selection on codon bias seems to have little influence over non-synonymous polymorphism or substitution patterns.

## Introduction

The genetic code is degenerate since 61 sense codons code for 20 amino acids, so most amino acids are encoded by more than one codon. These synonymous codons, however, are not equally used, a phenomenon known as codon usage bias (CUB) (Clarke 1970; Grantham, Gautier, Gouy, et al. 1980; Grantham, Gautier, and Gouy 1980; Post and Nomura 1980). The origins of such biases have been extensively debated. Evidence suggests that mutation bias, biased gene conversion, and natural selection influence CUB. However, the extent to which each factor affects codon usage depends on the species. While mutation bias and biased gene conversion seem to dominate in some species (Muto and Osawa 1987; Ohama et al. 1990; Wright and Bibb 1992), in others, such as *Escherichia coli*, selection plays a key role (Gouy and Gautier 1982; Sharp et al. 1993; Urrutia and Hurst 2001).

Various selective pressures have been proposed to affect synonymous sites, such as DNA stability, RNA stability, protein folding, gene regulation, and translational efficiency (Sharp and Li 1986; Eyre-walker and Bulmer 1993; Eyre-Walker 1996; Stoletzki and Eyre-Walker 2007; Plotkin and Kudla 2011). The latter is the most widely studied, with extensive evidence supporting this hypothesis (Akashi 1995; Eyre-Walker 1996; Stoletzki and Eyre-Walker 2007). One of the earliest observations of selection for translation efficiency was the correlation between the use of certain synonymous codons and tRNA concentrations (Ikemura 1985; Bulmer 1987). According to this theory, selection favours the use of more frequent codons, the so-called major codons, as these may increase translation efficiency. Such selective pressure is expected to be stronger in highly expressed or functionally constrained genes (Ikemura 1985; Sharp and Li 1986; Duret 2002), a pattern observed in *Drosophila melanogaster* (Akashi 1995) and *Escherichia coli* (Eyre-Walker 1996). Moreover, evidence suggests that genes with high codon bias have lower rates of synonymous substitutions than genes with low codon bias (Sharp and Matassi 1994; Plotkin and Kudla 2011). These observations suggest that selection acts upon codon usage during translation, a pattern observed in multiple organisms, including bacteria (P. M. Sharp and Li 1987; Berg and Martelius 1995), yeast (Bennetzen and Hall 1982; Sharp and Li 1986) *Drosophila* (Sharp and Li 1989), and nematodes (Stenico et al. 1994).

The component of translation that selection is acting upon remains the subject of some debate (Akashi and Eyre-Walker 1998; Reis et al. 2004; Stoletzki and Eyre-Walker 2007). The use of certain codons could potentially affect the elongation rate, the cost of proofreading or the accuracy of translation (Akashi and Eyre-Walker 1998; Reis et al. 2004; Stoletzki and Eyre-Walker 2007). However, as the initiation of translation is rate-limiting, selection on elongation is not to maximise the rate at which a particular gene is translated but to maximise the overall efficiency of translation by using ribosomes most efficiently. Hence, using optimal codons does not increase the expression of a gene, but it can increase the growth rate (Bulmer 1991; Weissman et al. 2021). Evidence of selection for translation accuracy is also widespread. In a ground-breaking study, Akashi (Akashi 1994) showed that the protein’s most important amino acid sites had a higher bias in *D. melanogaster*, an observation also made in *E. coli* (Stoletzki and Eyre-Walker 2007). Besides, the observed correlation between codon usage bias and gene length in *E. coli* (Eyre-Walker 1996) is also consistent with accuracy – it is more energetically expensive to make mistakes translating longer genes.

While the selective determinants of codon usage bias remain debated, one thing is clear: selection on codon usage depresses the rate of synonymous substitution in many species (Sharp and Li 1987; Sharp and Li 1989; Stenico et al. 1994; Berg and Martelius 1995). At the level of coding sequence evolution, however, little is known about the impact of codon usage bias. We hypothesise that selection for codon usage could also impede the rate of non-synonymous substitutions if two amino acids have different preferred codons. In this scenario, a non-synonymous mutation will be favoured by selection if it changes from a sub-optimal to an optimal codon, whereas the opposite will be disfavoured. For example, consider a non-synonymous mutation between Lysine (Lys) and Glutamine (Gln) in *E. coli*. As amino acids tend to use codons with high codon usage more frequently, we expect that a non-synonymous mutation changing Lys to Gln will generally involve a mutation from AAA, the preferred codon for Lys, to CAA, the unpreferred codon for Gln. One would therefore expect that selection on synonymous codon use will disfavour this mutation. Hence, selection on synonymous codon usage might reduce the level of non-synonymous polymorphism and the substitution rate between certain amino acids, those whose preferred codons differ (typically in the third codon position). Here, we investigate this theory in two bacterial species: *E*.*coli*, given the extensive evidence for selection on codon usage (Hershberg and Petrov 2008), and *Streptococcus pneumoniae*, as it has one of the strongest selective pressures on codon usage among bacteria (Sharp et al. 2005).

## Results

### Synonymous codon usage is a minor determinant of non-synonymous polymorphisms

We devised a series of tests to investigate whether selection on synonymous codon usage affects protein evolution. First, we assessed the impact of CUB on the segregating non-synonymous polymorphisms by considering all codon pairs separated by a single non-synonymous polymorphism (for example, AAA and CAA). Let us consider polymorphic sites for these two codons and infer the direction of mutation from the minor allele frequency. Thus, we infer an AAA>CAA mutation if CAA is the minor allele. Given that AAA is preferred for Lys in *E. coli* and CAA is unpreferred for Gln, we predict that the number of polymorphisms inferred to be CAA>AAA should be greater than AAA>CAA, assuming that there is no mutation bias or biased gene conversion and the average selective pressure in favour or against a Gln>Lys is the same as Lys>Gln (see Discussion). We can quantify the bias in favour of CAA>AAA in terms of an odds ratio, where we consider polymorphic sites inferred to be CAA>AAA, AAA>CAA, and monomorphic sites that are CAA and AAA (see File S1):

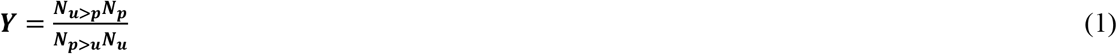

Where *N*_*u*>*p*_ and *N*_*p*>*u*_ are the numbers of unpreferred to preferred and preferred to unpreferred polymorphisms, respectively, where the unpreferred and preferred status are determined by each codon’s relative synonymous codon usage (RSCU, *i*.*e*., the observed frequency of each codon divided by the frequency expected when assuming equal usage for all synonymous codons within an amino acid (Sharp et al. 1986)); and *N*_*p*_ and *N*_*u*_ are the numbers of sites that are monomorphic for the preferred and unpreferred codons, respectively. We only consider pairs of codons where one codon’s RSCU value is above the average for its amino acid (RSCU > 1), and the other is below the average for the other amino acid (RSCU < 1). We sum values of *Y*, either for a codon pair across genes or for all codon pairs within a gene, using the Cochrane-Mantel-Haenzsel method (Mantel 1963) to avoid Simpson’s paradox, to yield a joint estimate of the OR - *Y*. As *Y* is a ratio of ratios, we present our results on the logarithmic scale. We used the RSCU values of codons amongst highly expressed genes, for which codon bias and selection are thought to be strongest. Positive *log(Y)* values mean that we have more non-synonymous polymorphisms changing from an unpreferred (low RSCU) to a preferred codon (high RSCU). We hypothesize that: (1) *log(Y)* should on average be greater than zero and (2) *log(Y)* should be correlated to the difference in the RSCU values between two codons.

We found little evidence for our first hypothesis in both species. The distribution of *log(Y)* values is symmetrically distributed around zero and the mean is not significantly different to zero both at the codon (Figure S1 and File S1; *E. coli:* mean = −0.0125, *p* = 0.9235, *S. pneumoniae*: mean = 0.0254, *p* = 0.8707) and gene levels (Figure S2 and File S1; *E. coli:* mean = −0.0287, *p* = 0.5557, *S. pneumoniae*: mean = −0.0952, *p* = 0.0475).

For our second hypothesis, we considered the value of *log(Y)* as a function of the differences in RSCU values between each pair of codons. For example, consider the codon pairs AAA<>CAA and AAA<>GAA. In *E. coli*, AAA is used 73% of the time in highly expressed genes, CAA 18%, and GAA 76%. Hence, we might expect AAA<>CAA to be affected by selection on synonymous codon use, but not AAA<>GAA, as the bias towards the use of AAA and GAA is very similar. Hence, we expect *Y* for AAA<>CAA to be greater than *Y* for AAA<>GAA. As expected, we found a significant positive correlation between *log(Y*) and the ratio of RSCU values of preferred to unpreferred codons (Figure 1 and File S1; *E. coli:* Pearson’s R = 0.31, *p* = 0.007; *S. pneumoniae:* R = 0.34, *p* = 0.006). We observed similar patterns when summing *Y* values across codon pairs within genes, where *log(Y)* is significantly positively correlated to the level of codon bias of the gene, as quantified by the codon adaptation index (CAI) (Sharp and Li 1987) (Figure 2 and File S1; *E. coli:* R = 0.12, *p* = 0.019; *S. pneumoniae:* R = 0.25, *p* = 4.9e-07). As expected, *log(Y)* is also significantly correlated to the level of gene expression (Figure S3 and File S1; *E. coli:* R = 0.16, *p* = 0.002; *S. pneumoniae:* R = 0.11, *p* = 0.03). These results suggest that selection influences the probability of observing an unpreferred to preferred polymorphism when the difference between their RSCU values is large. Given the overall distribution of *log(Y)* in both species, however, selection on synonymous codon usage appears to be a minor determinant of the level of non-synonymous polymorphism.

**Figure 1.**
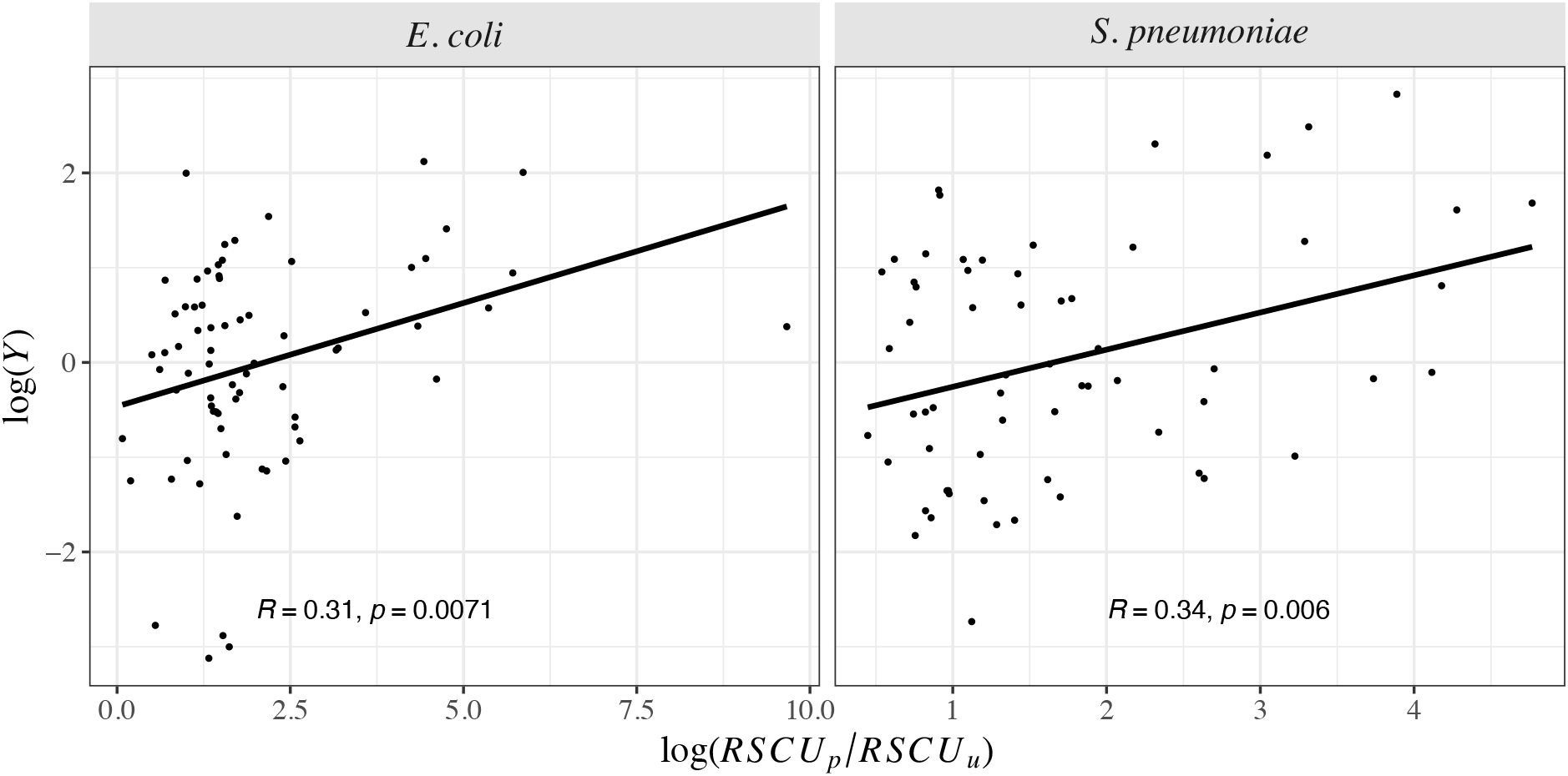
Relationship between the ratio of the RSCU value of preferred to unpreferred codons (log(*RSCU*_*p*_/*RSCU*_*u*_)) and *log(Y)* for *E. coli* (left) and *S. pneumoniae* (right). A linear model was fitted to the data and is represented with the dark line along with the Pearson correlation coefficient and the respective significance values.

**Figure 2.**
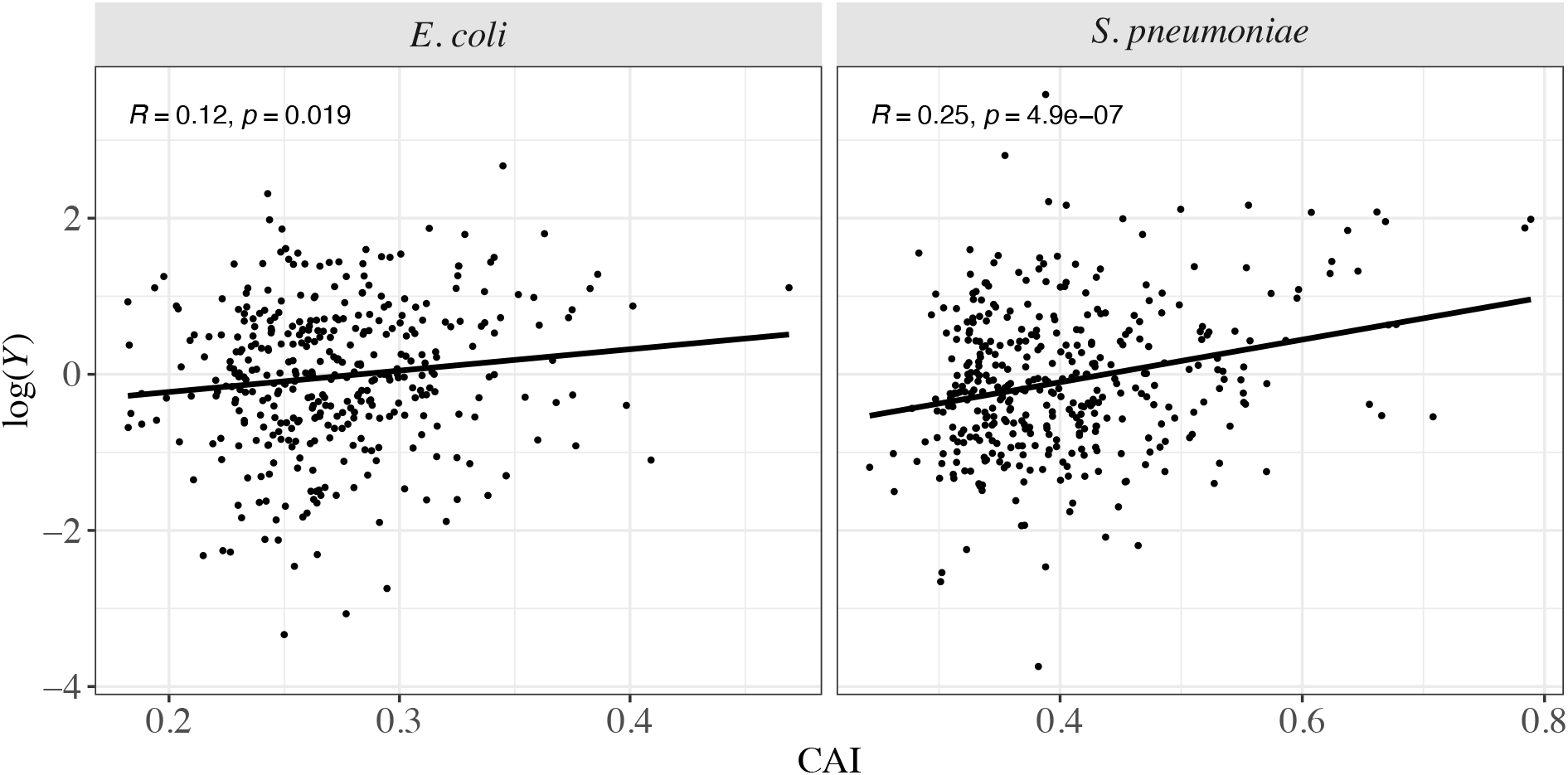
Relationship between *log(Y)* and the codon adaptation index (CAI) per gene. Legend as in Figure 1.

### The weak impact of codon usage bias on patterns of sequence divergence

We predict that selection on synonymous codon use slows the rate of non-synonymous substitution, a relationship that should be stronger between amino acids with different codon usage. To test this hypothesis, we quantified the average difference in codon usage between any two amino acids by estimating the expected change in log RSCU given a single hypothetical non-synonymous substitution assuming that all mutations are equally likely, a statistic we defined as *Z* (see Material and Methods and File S2). *Z* will take values around zero when two amino acids have similar codon usage patterns; positive when CUB positively impacts amino acid changes (*i*.*e*., an amino acid mutation increases the RSCU value on average); and negative if CUB impedes amino acid substitutions (*i*.*e*., an amino acid mutation reduces the RSCU value on average). We make three predictions: (1) *Z* should be skewed towards negative values given that a non-synonymous substitution will typically reduce the RSCU value (*i*.*e*., amino acids tend to use codons with high RSCU more frequently); (2) *Z* should be positively correlated with the number of substitutions between two amino acids (*d*_*ij*_); and (3) the relationship between *Z* and the rate of amino acid substitutions should become stronger with increasing codon usage (*i*.*e*., in highly expressed genes).

As expected, we found clear evidence for our first prediction, as *Z* is generally negative in both species (Figure S4 and File S2; *E. coli*: mean = −0.653, *S. pneumoniae*: mean = −0.464). By assessing the relationship between *Z* and the rate of sequence divergence for each amino acid pair (*d*_*ij*_), we observed a peak in *d*_*ij*_ values around 0, indicating that rates of sequence divergence are the highest between amino acids with similar codon usage preferences (Figure 3 and File S2). However, there is no correlation between *Z* and *d*_*ij*_ in the two species (Figure 3 and File S2; *E. coli*: R = −0.002, p = 0.99; *S. pneumoniae:* R = 0.13, p = 0.28). Finally, we hypothesized that the correlation between *Z* and *d*_*ij*_ should be stronger in highly expressed genes, as selection is thought to be more efficient. We assessed this relationship by analysing the correlation between Z and *d*_*ij*_ for each quartile of the distribution of CAI across genes. We found no differences in this relationship in both species, as there are no significant correlations between Z and *d*_*ij*_ in any of the CAI groups (Figure 4, Table S1 and File S2). However, this relationship seems to get stronger with lower CAI values, particularly in *S. pneumoniae* (Figure 4, Table S1 and File S2). These findings suggest that, although selection on codon usage impacts the probability that a non-synonymous substitution occurs, this relationship is merely weak.

**Figure 3.**
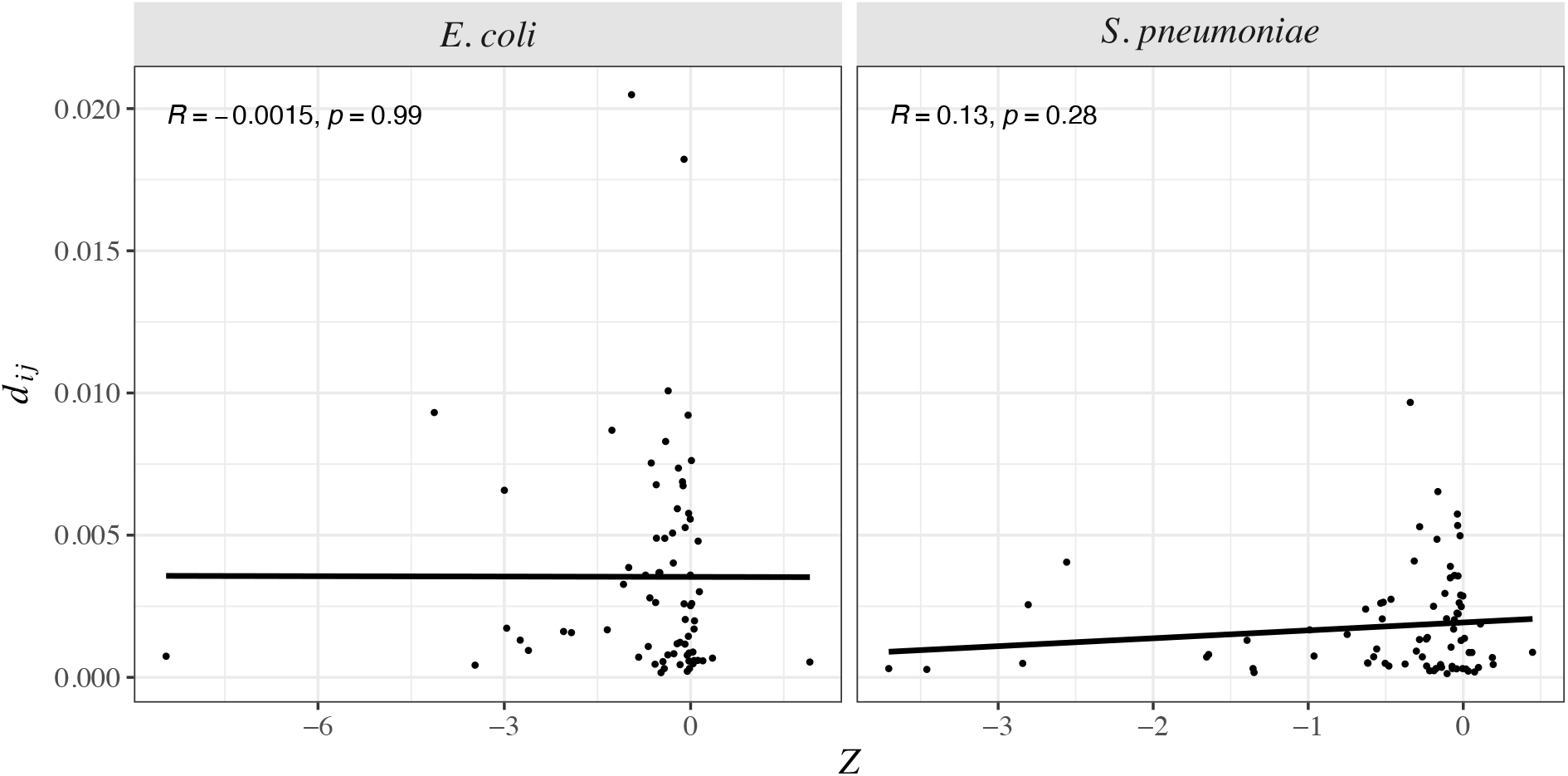
Relationship between sequence divergence (*d*_*ij*_) and *Z* for *E. coli* (left) and *S. pneumoniae* (right). Legend as in Figure 1.

**Figure 4.**
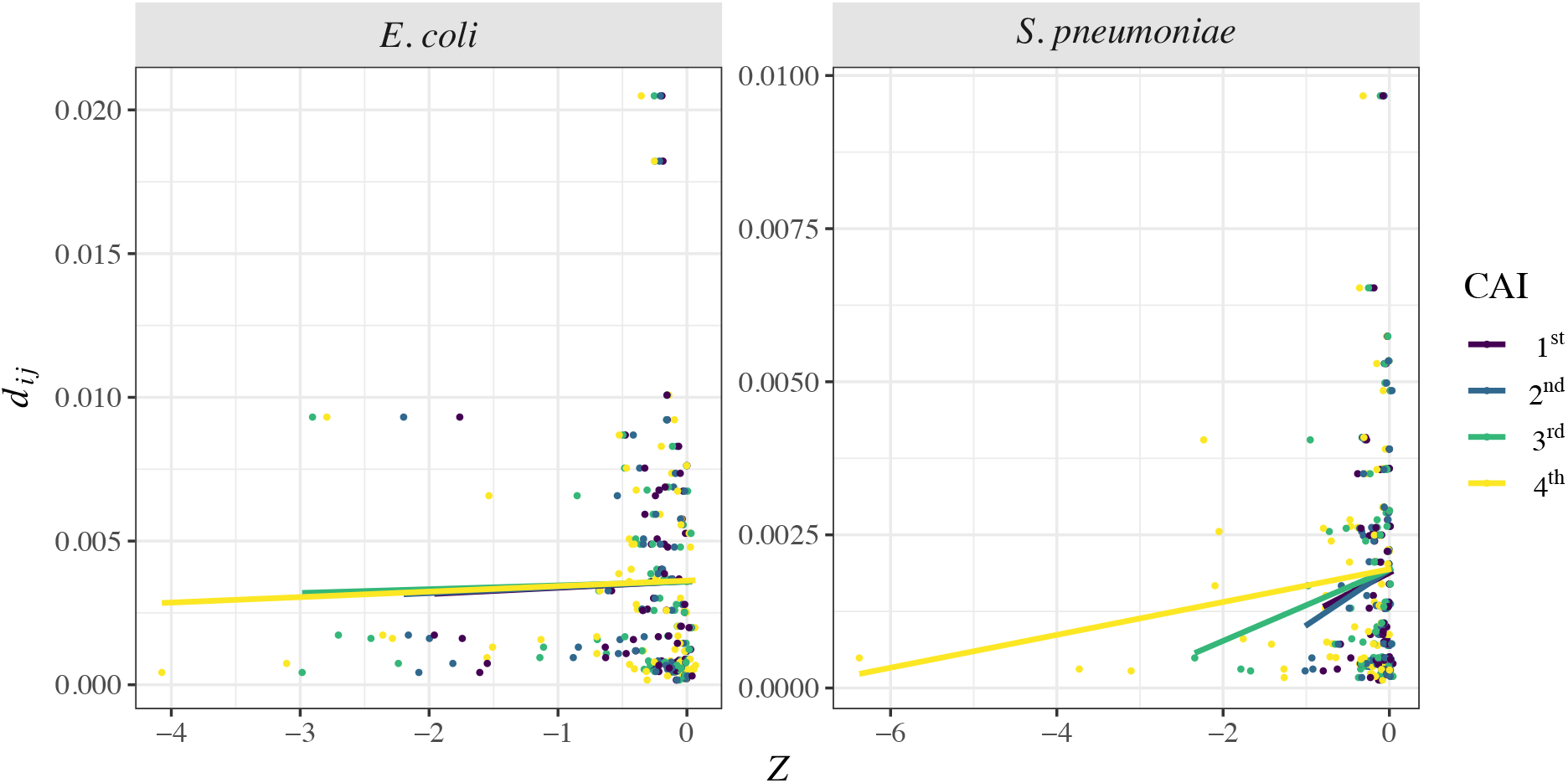
Relationship between Z and *d*_*ij*_ for different categories of CAI for *E. coli* (left) and *S. pneumoniae* (right). Each dot is coloured according to the respective CAI category. Legend as in Figure 2.

### The impact of mutation bias and biased gene conversion (BGC) on estimates of *log(Y)* and *Z*

As mutation bias and biased gene conversion are known to affect the composition of bacterial genomes (Hershberg and Petrov 2010; Hildebrand et al. 2010; Lassalle et al. 2015), we further assessed whether these could be biasing our results. We split our dataset into three groups: codon pairs that are not GC-biased, *i*.*e*., not affected by biased gene conversion (no-bias: A<>T and G<>C mutations), codon pairs where A or T are the selectively preferred alleles (pAT), and codon pairs where G or C are the selectively preferred alleles (pGC). When looking at the distribution of *log(Y)*, we observed a clear difference between mutation categories, with the average *log(Y)* for pAT mutations being significantly positive in both species (Figure S5 and File S3; *E. coli:* mean = 0.879, *p* = 2.45e-05; *S. pneumoniae*: mean = 1.301, *p* = 2.61e-07), while being significantly negative for pGC mutations (Figure S5 and File S3; *E. coli:* mean = −0.795, *p* = 4.64e-04; *S. pneumoniae*: mean = −1.016, *p* = 7.23e-08). For the no-bias mutations, the distribution of *log(Y)* is centred around zero, being only significantly positive in *E. coli* (Figure S5 and File S3; *E. coli:* mean = 0.297, *p* = 2.509e-02; *S. pneumoniae*: mean = −0.07, *p* = 0.722). These results suggest that there are more polymorphisms changing from unpreferred to preferred codon when A or T are the preferred allele, thus supporting the overall bias towards AT mutations previously observed in bacteria (Hershberg and Petrov 2010; Hildebrand et al. 2010). By analysing the relationship between *log(Y)* and *log(RSCU*_*p*_*/RSCU*_*u*_), we observed that it is still positive for all mutation types, although being weaker in *E. coli* and no-bias codons in *S. pneumoniae* (Figure S6 and File S3). Nonetheless, by doing an ANCOVA, we find the relationship between *log(Y)* and *log(RSCU*_*p*_*/RSCU*_*u*_) is significant in both species (*E. coli*: *p* = 0.001; *S. pneumoniae*: *p* = 1.437e-05) (Figure S6 and File S3). Hence, despite the clear mutation bias towards AT mutations, this does not seem to impact our overall result: there are more polymorphisms changing from unpreferred to preferred codons when their differences in RSCU are larger, a pattern that is independent of the mutation type.

To further assess the impact of selection on codon usage among mutation categories, we estimated the mean differences in allele frequencies between the preferred and unpreferred alleles for each codon pair (*i*.*e*., mean allele frequency of the preferred allele - mean allele frequency of the unpreferred allele; see File S3). Our results showed that the average differences in allele frequencies tend to be positive in no-bias and pGC codon pairs and negative in pAT mutations, being significantly different only in *E. coli* (ANOVA *E. coli: p* = 0.008; *S. pneumoniae: p =* 0.447; Figure S7 and File S3). When looking at the relationship between *log(RSCU*_*p*_*/RSCU*_*u*_) and differences in allele frequencies, we observed a positive correlation for all mutation categories in *E. coli*, being significant in no-bias and pGC codon pairs (Figure S8 and File S3). In *S. pneumoniae*, however, this pattern only holds for pGC mutations (Figure S8 and File S3). These findings agree with the BGC theory, as GC polymorphisms segregate at higher frequencies than AT polymorphisms. Hence, a mutation changing from an unpreferred to a preferred allele is more likely to increase the mean allele frequency if it occurs in the direction of AT>GC.

At the substitution level, the type of mutation had no impact on our results. The distribution of *Z* is skewed towards negative values (Figure S9 and File S3) and the correlation between Z and *d*_*ij*_ continues to be weak across all mutation types (Figure S10 and File S3).

## Discussion

We predict that selection on synonymous codon use should affect the level of non-synonymous polymorphism and substitution. However, we find little evidence of this both at the polymorphism and divergence levels. We do not find that non-synonymous polymorphisms changing from an unpreferred to a preferred codon are not more prevalent than the opposite (Figure S1). And we do not find that rates of non-synonymous substitution are lower between amino acids where the average change in RSCU is predicted to be greater (Figure 3). However, we do find that unpreferred to preferred polymorphisms amongst pairs of codons that differ substantially in their RSCU values are more frequently observed than the opposite and that these mutations tend to segregate at higher frequencies (Figures 1 and 2, Figure S8). This pattern may explain the other results. We only observe an effect of codon usage on non-synonymous polymorphisms when the differences between codons are substantial. However, such codon pairs are relatively rare. Most non-synonymous mutations do not change the RSCU value very substantially (average change in RSCU = 1.899). Hence, it is perhaps not surprising that we do not observe any effect of selection on codon usage on patterns of protein evolution. Moreover, although we observed an impact of mutation bias (Figure S5) and BGC (Figure S7) on the polymorphism data, these do not seem to impact our overall patterns of selection on codon usage at the polymorphism (Figure S6) and substitution levels (Figure S10).

However, there may be other reasons for this lack of effect. On the one hand, we are assuming that if two codons from two different amino acids have the same RSCU value, they are equally fit in terms of translational efficiency. Moreover, if codon X has a higher RSCU value than codon Y, it is more fit. However, this is unlikely to be true due to different binding affinities between codons and tRNAs and the absolute tRNA concentrations. All codons for one amino acid may be fitter than those for another amino acid. This would add noise to our analysis, making it harder to detect an effect.

On the other hand, our model assumes directional selection in favour of codon usage, where unpreferred codons exist primarily because the system is in mutation-selection balance, *i*.*e*., selection is sufficiently weak in favour of the preferred codons that not all sites are occupied by them (Bulmer 1991). However, there are two alternative theories for why unpreferred codons exist. The first theory states that some unpreferred codons are used because the same sequence can have different functions, a phenomenon known as antagonistic pleiotropy (Akashi and Eyre-Walker 1998). Such phenomenon was well-characterized in *E. coli*, where the weaker bias in codon usage at the start of the genes was attributed to selection against secondary structure in the mRNA (Eyre-walker and Bulmer 1993; Kudla et al. 2009). If antagonistic pleiotropy is prevalent, a low RSCU codon might be fitter at a particular site than a high RSCU codon. So, although high RSCU codons tend to be fitter than low RSCU codons, considerable deviations from this norm might obscure the patterns we predict. Second, codon usage might be subject to stabilizing rather than directional selection. Indeed, experiments with yeast have demonstrated that the average time for the ribosome to move through a codon is not correlated to the RSCU value or the concentration of the tRNA matching the codon (Qian et al. 2012). If selection is stabilizing, changing a preferred to unpreferred codon is equally strongly selected as the opposite. However, our observation that *log(Y)* is positively correlated to the ratio of the RSCU values is more consistent with the directional selection model.

Intriguingly, we find a stronger influence of the mutation type on patterns of polymorphism when compared to selection on codon usage. *Log(Y)* tends to be positive for pAT codon pairs, negative for pGC codon pairs, and, depending on the species, either positive or negative for no-bias mutations (Figure S5). This pattern supports the known AT-biased mutation pattern in *E. coli* (Lee et al. 2012) and the prevailing AT bias in most bacteria (Hershberg and Petrov 2010; Hildebrand et al. 2010). When considering allele frequencies, we observed a pattern consistent with GC-biased biased gene conversion, as pGC codon pairs segregate at higher frequencies than pAT codon pairs (Figure S7).

Overall, we found little evidence for the impact of codon usage on the rates of protein evolution in bacteria, being only evident when the codons differ substantially in their codon usage. Moreover, we observed that the relationship between codon bias and rates of non-synonymous substitution varies between species, being stronger in *S. pneumoniae*. This pattern might reflect differences in the strength of selection acting at synonymous sites, as Sharp et al. (Sharp et al. 2005) found relatively higher selection coefficients in *S. pneumoniae*. Our study, therefore, provides evidence for a subtle impact of codon usage bias on rates of protein evolution, a relationship that may depend on the strength of selection acting at the synonymous level.

## Material and Methods

### Population genomic data and data filtering

Polymorphism data was downloaded from the ATGC database (Kristensen et al. 2017), where full orthologous genome alignments were obtained for *Escherichia coli* (ATGC001) and *Streptococcus pneumoniae* (ATGC003). We filtered the data to keep only genes present in the core genome, resulting in a total of 1,316 and 1,194 genes in 164 and 31 strains of *E. coli* and *S. pneumoniae*, respectively. These alignments were then split by gene and codon statistics were obtained with the BppPopStats program (Guéguen et al. 2013).

### Divergence analysis

Divergence was estimated by taking one outgroup species belonging to the same ortholog group of each organism: *Salmonella enterica* and *S. mitis* for *E. coli* and *S. pneumoniae*, respectively. Pairwise alignments for each species pair were obtained by taking the longest sequence for each gene, resulting in a total of 1,291 and 1,123 genes for analysis. The genetic divergence between amino acid *i* and *j* for each species pair (*d*_*ij*_), was estimated for all possible amino acid pairs with the following equation:

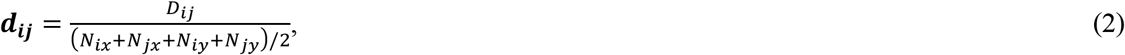

Where *D*_*ij*_ is the number of codon sites at which species *x* has amino acid *i* and species *y* has *j* and *vice versa*, and *N*_*ix*_ is the number of sites at which species *x* has amino acid, and *N*_*iy*_ is the number of sites at which species y has amino acid j.

### Gene expression

For *E. coli*, we used the compiled gene expression data set from the DREAM5 network inference challenge (Marbach et al. 2012), resulting in a total of 1,287 genes for analysis. For *S. pneumoniae*, gene expression data were obtained from the Gene Expression Omnibus (GEO) database (Edgar et al. 2002) where we used the normalised data set GSE77587, ending up with 803 genes for further analysis.

### Correspondence analysis and RSCU estimates

Correspondence analyses were performed on the two species using the strains *E. coli K12 MG1655* and *S. pneumoniae R6*. The absolute frequency of codons (AF) and the relative synonymous codon usage (RSCU) were estimated using the function “uco” from the “seqinr” R package (Charif and Lobry 2007) for the full set of genes. The correspondence analyses (CA) were performed with the function “dudi.coa” from the “ade4” R package (Dray and Dufour 2007) using the RSCU values (CA-RSCU) and the AF (CA-AF). The within-group correspondence analyses (WCA) were performed using CA-AF (WCA-CA-AF). We compared the two methods (CA-RSCU and WCA-CA-AF) to check whether there were significant differences between them. The first two principal components were used to assess the amount of variation in codon usage bias across genes. We identified genes showing a greater bias in synonymous codon usage by looking at the distribution of ribosomal proteins and highly expressed genes on PC1. RSCU values of highly expressed genes were then estimated for the set of genes belonging to the first quantile of this distribution (*i*.*e*., the top 10%). The relative adaptiveness of each codon (*w*) and the codon adaptation index (CAI) were estimated using these RSCU values as in Sharp & Li (Paul M. Sharp and Li 1987).

### The *log(Y)* statistic

At the polymorphism level, we filtered the data to keep only non-synonymous codon pairs separated by one non-synonymous mutation to avoid any further mutational bias. We only considered codon pairs in which the codon changes from unpreferred (defined as a codon having an RSCU value below one) to a preferred codon (defined as having an RSCU value above one). The final data sets included 72 and 63 codon pairs for *E. coli* and *S. pneumoniae*, respectively. To assess the impact of codon usage bias on patterns of non-synonymous polymorphisms, we developed a statistic we denote as *Y*. This statistic is based on the Cochrane-Mantel-Haenzsel method (Mantel 1963) for combining contingency tables. With this test, we ask whether the ratio of polymorphic to non-polymorphic sites is the same for preferred (p) and unpreferred (u) codon sites. Our contingency tables are composed of: (*N* _*u*>*p*_) polymorphic u>p sites; (*N*_*p*>*u*_) polymorphic p>u sites; (*N*_*u*_) non-polymorphic u sites; and (*N*_*p*_) non-polymorphic p sites. The odds ratio is estimated as in equation 1 and the joint odds-ratio (OR, here defined as *Y*) is then estimated by:

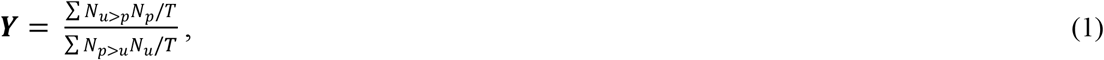

where T = *N*_*u*>*p*_ + *N*_*p*>*u*_ + *N*_*u*_ + *N*_*p*_. This sum can be applied for each codon pair across genes or for each gene across codon pairs. Our results are present in terms of *log(Y)*, where positive values mean that there are more non-synonymous polymorphisms between codons that change from low to high RSCU values (see detailed R scripts in File S1).

This method, however, may be biased in the way the numbers of polymorphic u and p sites are counted because we have excluded all codons that do not segregate the polymorphism of interest (e.g., GAA<>AAA, as these are both preferred codons in *E. coli*) or that are fixed in the species for AAA or GAA. As there might be synonymous and other non-synonymous polymorphisms also segregating within our preferred and unpreferred codons, we will be removing some sites from consideration. This is expected to differentially affect the preferred and non-preferred codons differently. For example, we expect synonymous preferred to unpreferred polymorphisms to be more common than the opposite. Consequently, we will tend to remove more preferred codons from our analysis, particularly those sites which might have otherwise been considered to be fixed for a codon. This will tend to result in *Y* > 1. To correct this potential bias, we altered the *Y* method to consider all sites for which the focal codon is the most common allele as monomorphic. So, in this contingency table we have: (*N*_*u*>*p*_) polymorphic u>p sites; (*N*_*p*>*u*_) polymorphic p>u sites; (*N*_*u*_) non-polymorphic u sites; (*N*_*p*_) non-polymorphic p sites; (*N*_*u*+_) polymorphic and u the most frequent allele; and (*N*_*p*+_) polymorphic and p the most frequent allele. The second joint odds-ratio (*Y’*) is then estimated by:

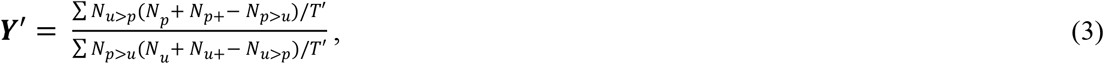

where T’ = *N*_*u*>*p*_ + *N*_*p*>*u*_ + (*N*_*u*_ + *N*_*u*+_ - *N*_*u*>*p*_) + (*N*_*p*_ + *N*_*p*+_ - *N*_*p*>*u*_). We subtract *N*_*u*>*p*_ and *N*_*p*>*u*_ to new measures of monomorphic sites because these are the sites for which u and p alleles are the most frequent in the population (see detailed R scripts in File S1).

### The *Z* statistic

At the divergence level, we applied the same filtering step to keep only non-synonymous codon pairs separated by one non-synonymous mutation. We assessed the impact of codon usage bias on the rate of protein divergence through a statistic we defined as *Z*. With this statistic, we are measuring the average change in RSCU given a single mutation. We do this by estimating the average counts of each codon in each species weighted by the logarithmic scale of the ratio of the RSCU value of codon *k* in amino acid *i* (*RSCU*_*ik*_) to the RSCU value of codon *g* in amino acid *j* (*RSCU*_*ig*_) (log (*RSCU*_*ik*_/*RSCU*_*ig*_):

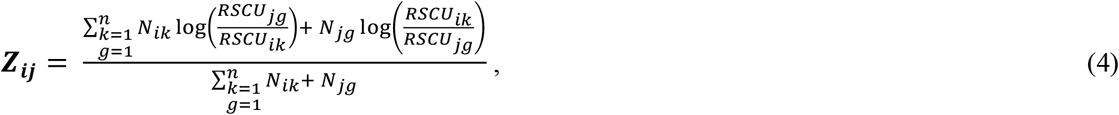

where *N*_*ik*_ = counts of codon *k* in amino acid *i*; *N*_*jg*_ = counts of codon *g* in amino acid *j*; and *n* is the total number of codon pairs for each amino acid pair. Negative values of this statistic mean that amino acid changes are impeded by codon usage bias (see detailed R scripts in File S2).

To analyse the relationship between *Z* and *d*_*ij*_ across CAI categories, we used the RSCU values estimated using the genes belonging to the respective CAI category to better assess the strength of selection acting on each group of genes (see detailed R scripts in File S2).

### Biased gene conversion (BGC) and mutation bias

To assess the impact of BGC and mutation bias, we subset our data sets into three categories of mutations: (1) codon pairs not affected by BGC (no-bias: A<>T and G<>C mutations); (2) codon pairs affected by BGC where A or T are the preferred alleles (pAT: GC<>AT where GC are unpreferred and AT preferred); and (3) codon pairs affected by BGC where G or C are the preferred alleles (pGC: GC<>AT where AT are unpreferred and GC preferred). This analysis was performed both at the polymorphism and divergence levels (see detailed R scripts in File S3). Differences in allele frequency were estimated using the mean minor allele frequency at each codon site, *i*.*e*., taking the folded site frequency spectrum (SFS; see File S3). As some sites were not successfully sampled in all individuals, we randomly downsampled the data to include 71 and 15 SFS categories for *E. coli* and *S. pneumoniae*, respectively. Differences in allele frequencies were estimated between codon pairs changing between unpreferred and preferred codons by taking the mean allele frequency of the preferred codon and subtracting the mean allele frequency of the unpreferred codon (see detailed R scripts in File S3).

### Statistical analysis

Statistical significance was assessed with Pearson’s correlation coefficient. All figures were plotted using the R package “ggplot2” (Wickham 2017). Statistical significance for the analysis of the type of mutation was assessed with an analysis of covariance (ANCOVA). A T-test was used to assess whether the means of the distribution of *log(Y)* were significantly different from zero.

## Acknowledgements

The authors thank Andrea Betancourt for the fruitful discussions. AEW thanks GBE for the funding.

