## Supplementary Data for "The silent impact: codon usage bias and protein evolution in bacteria"

**Supplementary Tables**

**Table S1.** Pearson's correlation coefficients between $Z$ and $d_{ij}$ for each category of CAI.

| | ***CAI*** | ***R*** | ***p-value*** |
|---|---|---|---|
| ***E. coli*** | 1st | 0.025 | 0.838 |
| | 2nd | 0.028 | 0.819 |
| | 3rd | 0.023 | 0.849 |
| | 4th | 0.039 | 0.747 |
| ***S. pneumoniae*** | 1st | 0.062 | 0.593 |
| | 2nd | 0.109 | 0.347 |
| | 3rd | 0.132 | 0.252 |
| | 4th | 0.145 | 0.207 |

**Supplementary Figures**

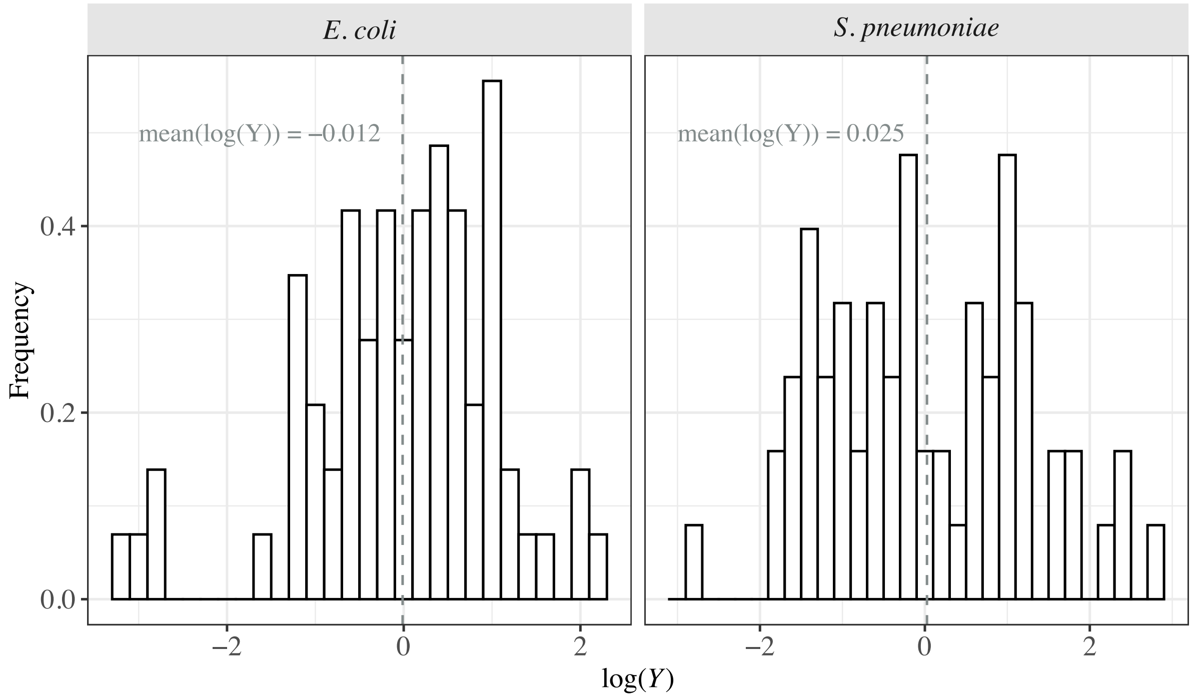


**Figure S1.** Distribution of *log(Y)* across codon pairs. The dashed line represents the mean value of the distribution. The binning size was set at 0.2.

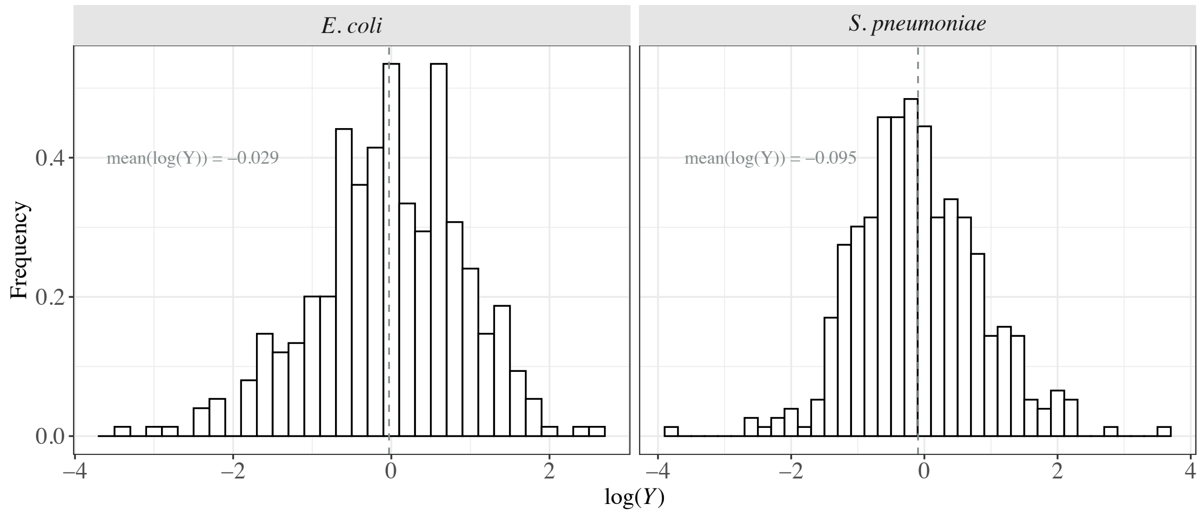


**Figure S2.** Distribution of *log(Y)* across genes. Legend as Figure S1.

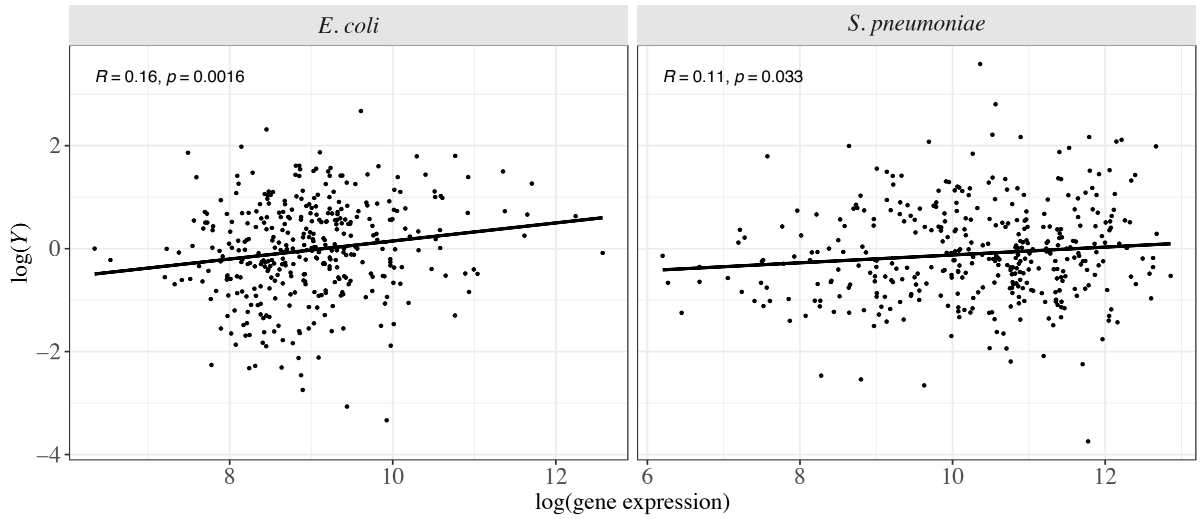


**Figure S3.** Relationship between *log(Y)* and mean gene expression levels for *E. coli* (left) and *S. pneumoniae* (right). A linear model was fitted to the data and is represented with the dark line along with the Pearson correlation coefficient and the respective significance values.

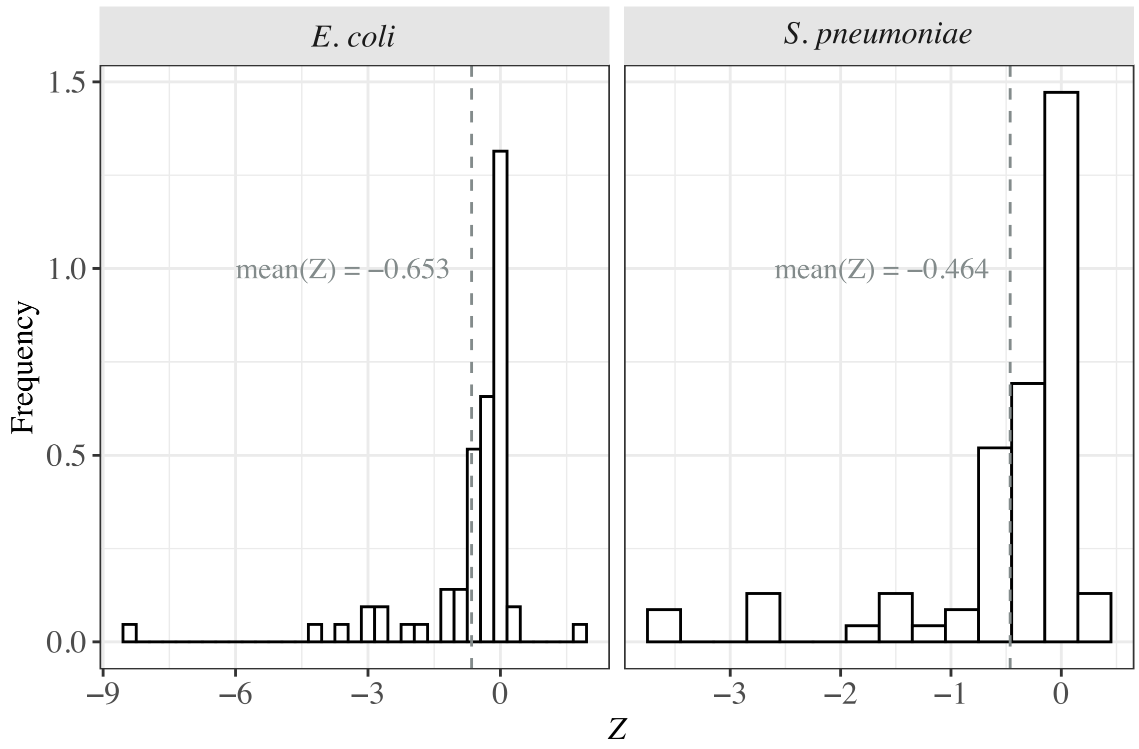


**Figure S4.** Distribution of *Z* across amino acid pairs for *E. coli* (left) and *S. pneumoniae* (right). The mean value of *Z* is represented by the dashed grey line. The binning size was set at 0.3.

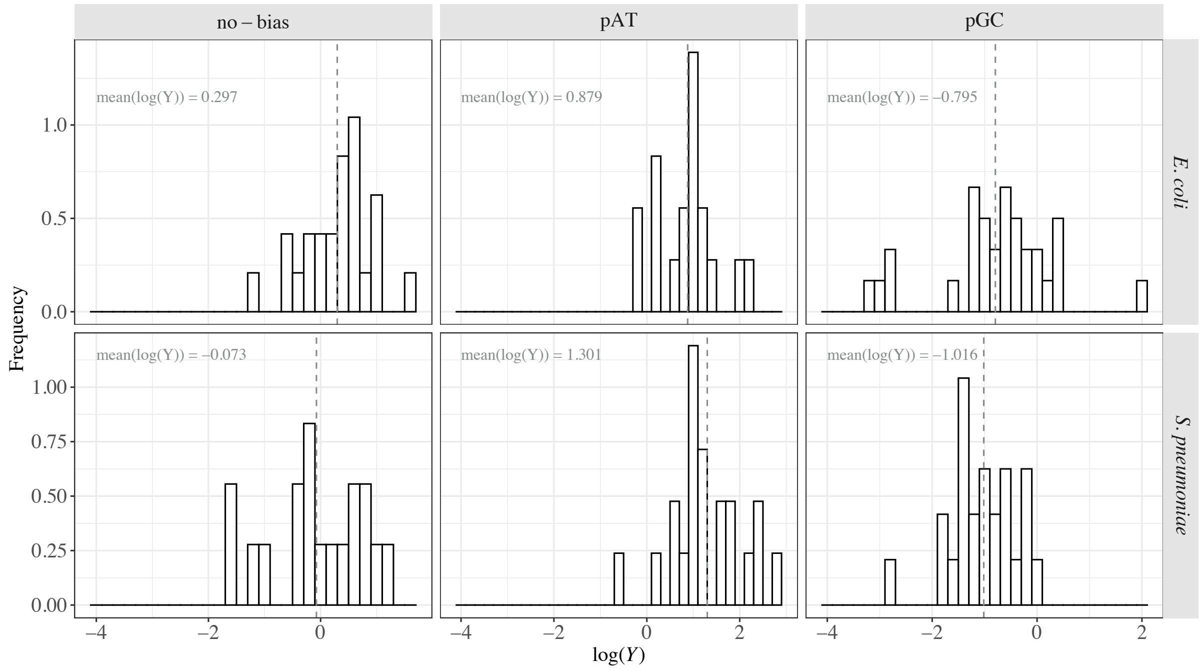


**Figure S5.** Distribution of *log(Y)* across codon pairs for each mutation type (no-bias = A<>T and G<>C mutations; pAT = mutation where A/T are the preferred alleles; pGC = mutation where G/C are the preferred alleles) in *E. coli* (top) and *S. pneumoniae* (bottom). The dashed line represents the mean value of the distribution. The binning size was set at 0.2.

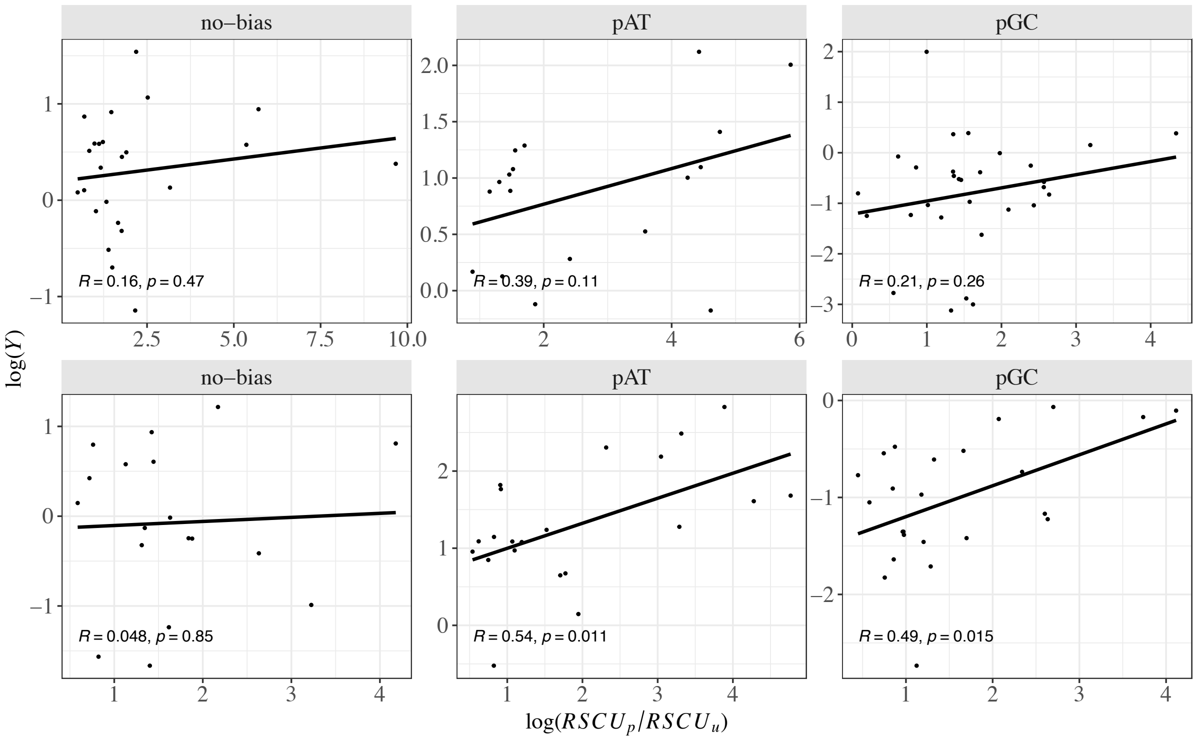


**Figure S6.** Relationship between the ratio of the RSCU value of preferred to unpreferred codons (log($RSCU_p/RSCU_u$)) and *log(Y)* for each mutation type (see legend of Figure S5) in *E. coli* (top) and *S. pneumoniae* (bottom). A linear model was fitted to the data and is represented with the dark line along with the Pearson correlation coefficient and the respective significance values.

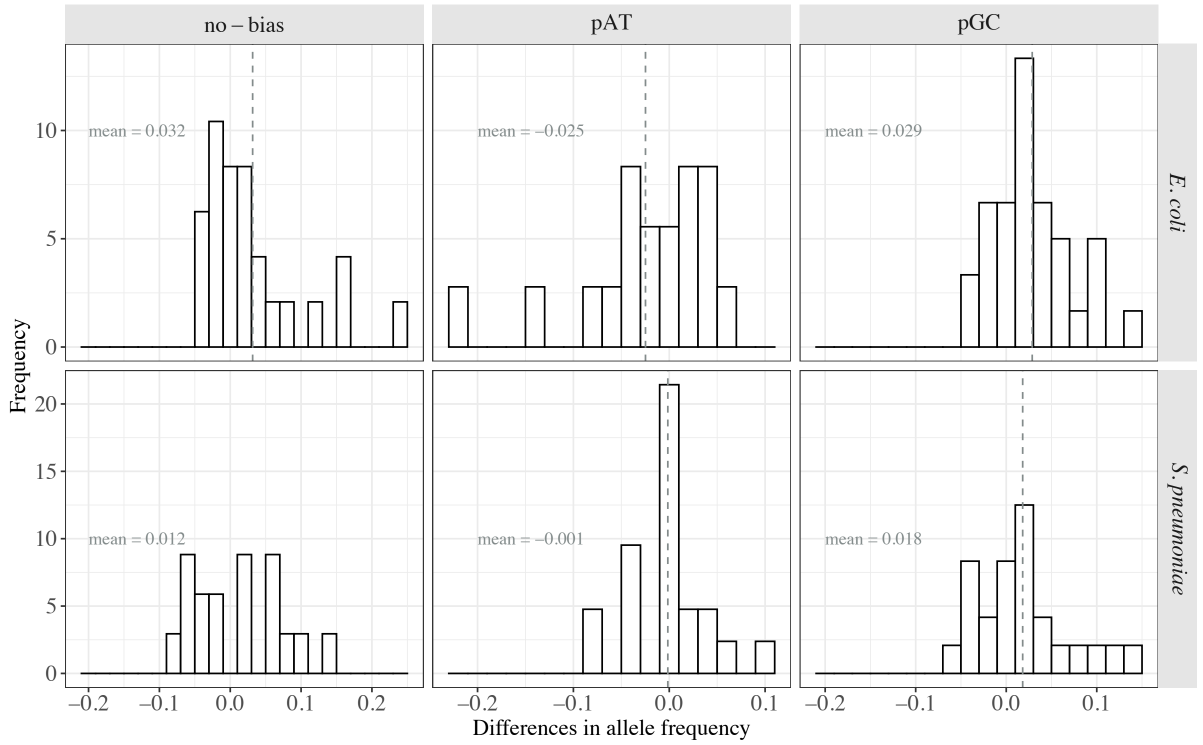

**Figure S7.** Distribution of the mean differences in allele frequency for each codon pair across genes for each mutation type (see legend of Figure S5) in *E. coli* (top) and *S. pneumoniae* (bottom). The dashed line represents the mean value of the distribution. The binning size was set at 0.02.

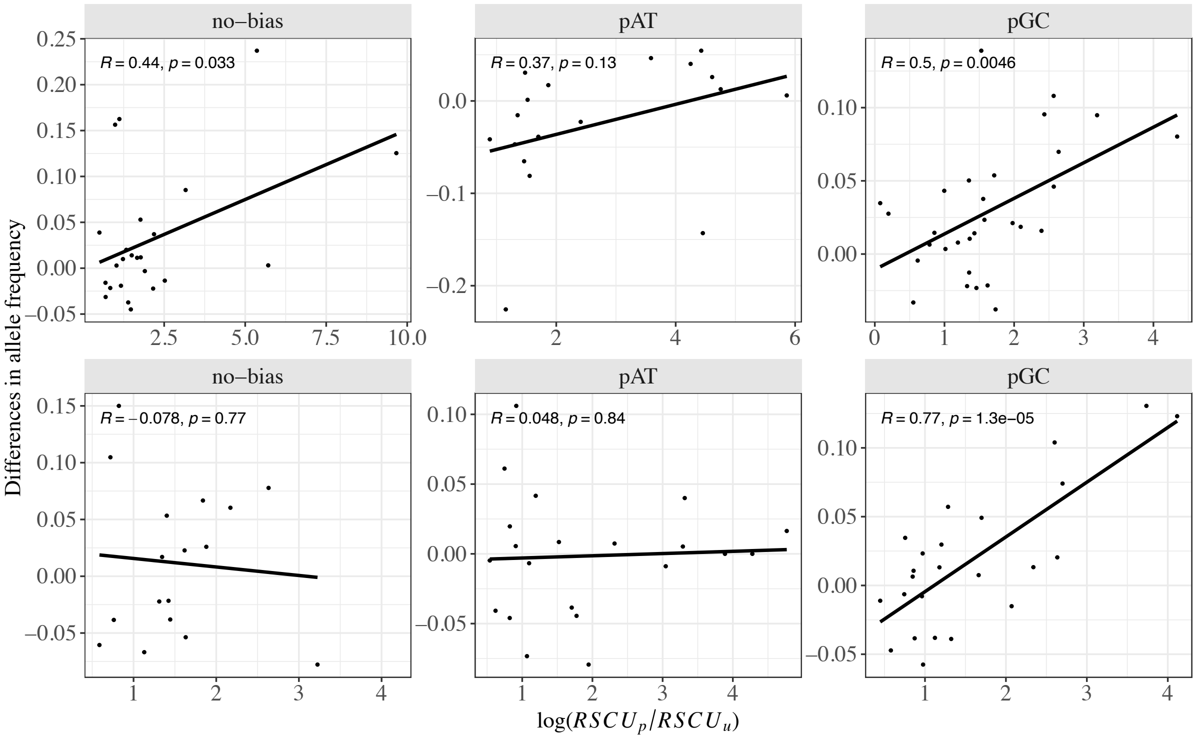


**Figure S8.** Relationship between the ratio of the RSCU value of preferred to unpreferred codons ($\log(RSCU_p/RSCU_u)$) and mean differences in allele frequency for each mutation type (see legend of Figure S5) in *E. coli* (top) and *S. pneumoniae* (bottom). A linear model was fitted to the data and is represented with the dark line along with the Pearson correlation coefficient and the respective significance values.

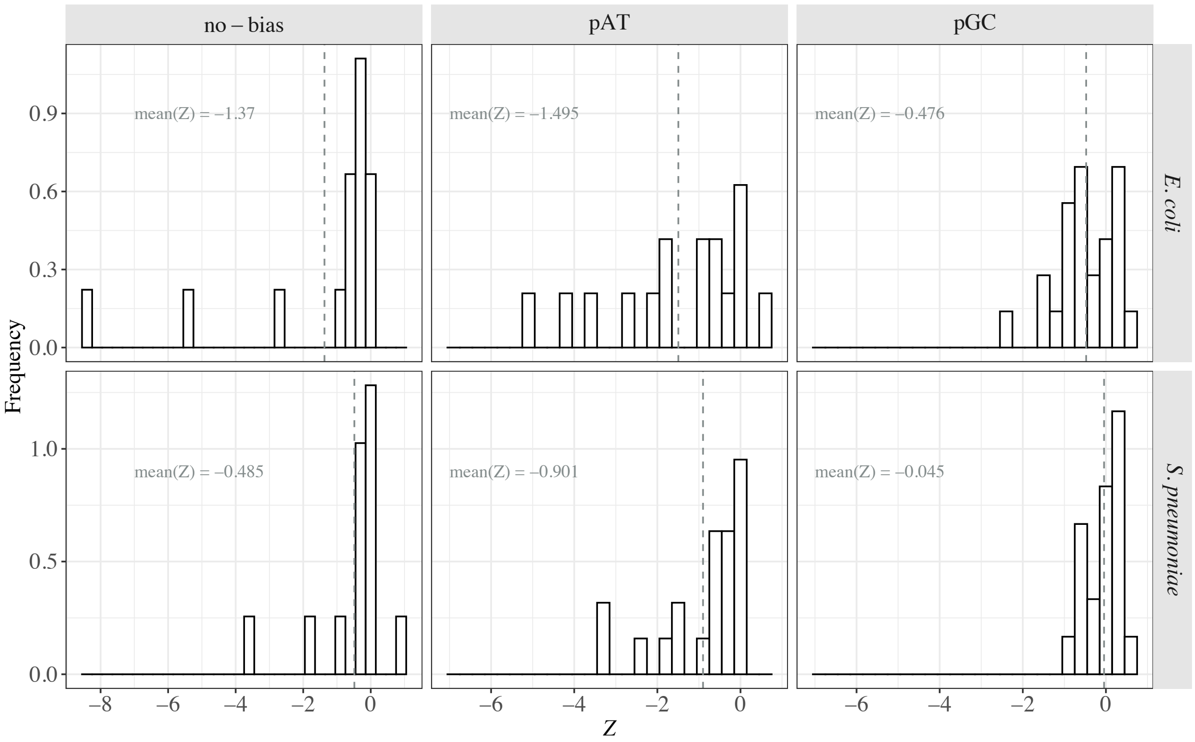


**Figure S9.** Distribution of Z across codon pairs for each mutation type (see legend of Figure S5) in *E. coli* (top) and *S. pneumoniae* (bottom). The dashed line represents the mean value of the distribution. The binning size was set at 0.3.

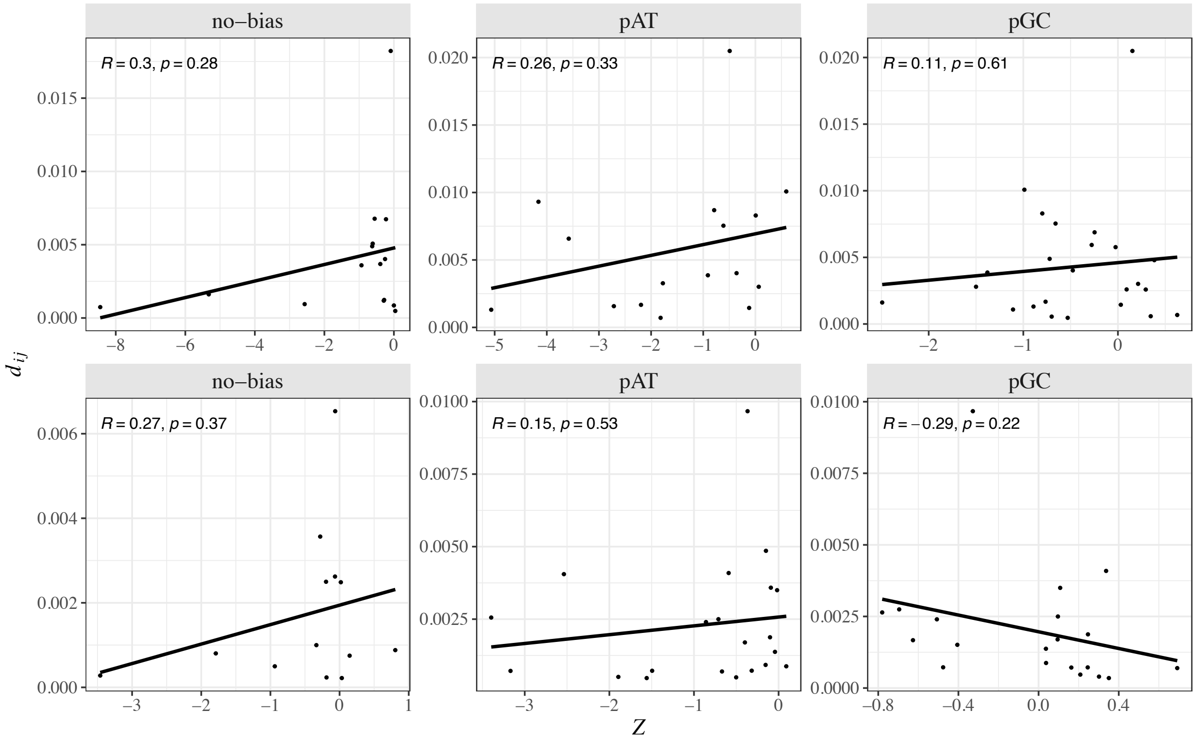


**Figure S10.** Relationship between sequence divergence ($d_{ij}$) and $Z$ in *E. coli* (top) and *S. pneumoniae* (bottom) for the different mutation types (see legend of Figure S5).
