## Supplementary material for "The silent impact: codon usage bias and protein evolution in bacteria": File S1

### Polymorphism analyses

af_moutinho

2022-07-04

This RMarkdown contains all analyses performed at the polymorphism level. The first chunk of the script reproduces the estimation of log(Y) in *E. coli* and *S. pneumoniae*.

```
# Libraries
library(data.table)
library(dplyr)
library(plyr)
library(ggplot2)
library(ggpubr)
library(kableExtra)
#

# getting the table with the data for each codon site:
codons_poly <- fread(file = "~/Dropbox/CUB_supplementary_data/tables/polymorphism_codons_tbl.csv",
                     sep = "\t", header = T)

# keeping only monomorphic and biallelic sites for analysis:
codons_poly2 <- codons_poly[codons_poly$NbAlleles == 2 | codons_poly$NbAlleles == 1,]


### calculations for the contingency tables:

# keeping only alleles with a single nucleotide difference
single_codon_pair <- codons_poly2[codons_poly2$nt.dif == 1,]

# counting the number of monomorphis p and u sites:
AB_counts <- ddply(single_codon_pair, c("species", "coID", "aa3_pair", "aa_minor",
                                        "aa_major", "codon_pair", "MajorAllele",
                                        "MinorAllele", "major_codon", "minor_codon"),
                   function(x) {
                              # A sites (np>p non-synonymous)
                              A <- sum(nrow(x[x$major_codon == "np" & x$minor_codon == "p"
                                              & x$IsSynPoly == 0 & x$NbAlleles == 2,]))
                              # B sites (p>np non-synonymous)
                              B <- sum(nrow(x[x$major_codon == "p" & x$minor_codon == "np"
                                              & x$IsSynPoly == 0 & x$NbAlleles == 2,]))
                              data.frame(A, B)
                            })


# counting the numbers of polymorphic sites:
polymorphic_counts <- ddply(codons_poly2, c("species", "coID", "MajorAllele"),
```

```
                             function(x) {
  # all polymorphic sites for which the major allele is unpreferred (poly_u_major)
  E <- sum(nrow(x[x$major_codon == "np" & x$NbAlleles == 2,]))
  # all polymorphic sites for which the major allele is preferred (poly_p_major)
  G <- sum(nrow(x[x$major_codon == "p" & x$NbAlleles == 2,]))
  C <- sum(nrow(x[x$NbAlleles == 1 & x$major_codon == "np",]))
  D <- sum(nrow(x[x$NbAlleles == 1 & x$major_codon == "p",]))
  data.frame(E, G, C, D)
})


# combining the tables:
codon_counts <- merge(AB_counts, polymorphic_counts,
                      by = c("coID", "MajorAllele", "species"))

colnames(codon_counts)[c(15:16)] <- c("C1", "D1")

# will now combine with the minor allele for counts of C and D:
sub_poly_counts <- subset(polymorphic_counts, select = -c(E, G))
codon_counts <- merge(codon_counts, sub_poly_counts,
                      by.x = c("coID", "MinorAllele", "species"),
                      by.y <- c("coID", "MajorAllele", "species"))

colnames(codon_counts)[c(17:18)] <- c("C2", "D2")

codon_counts$C <- codon_counts$C1 + codon_counts$C2
codon_counts$D <- codon_counts$D1 + codon_counts$D2

codon_counts_df <- subset(codon_counts, select = - c(C1, D1, C2, D2))

# making the counts of each cross:
codon_counts_df$sum <- codon_counts_df$A + codon_counts_df$B + codon_counts_df$C +
  codon_counts_df$D
codon_counts_df$C2 <- codon_counts_df$C + codon_counts_df$E - codon_counts_df$A
codon_counts_df$D2 <- codon_counts_df$D + codon_counts_df$G - codon_counts_df$B
codon_counts_df$sum2 <- codon_counts_df$A + codon_counts_df$B + codon_counts_df$C2 +
  codon_counts_df$D2
codon_counts_df$cross_p <- (codon_counts_df$A*codon_counts_df$D)/codon_counts_df$sum
codon_counts_df$cross_u <- (codon_counts_df$B*codon_counts_df$C)/codon_counts_df$sum
codon_counts_df$cross_p2 <- (codon_counts_df$A*codon_counts_df$D2)/codon_counts_df$sum2
codon_counts_df$cross_u2 <- (codon_counts_df$B*codon_counts_df$C2)/codon_counts_df$sum2

# keeping only non-synonymous polymorphisms and the ones that lead to a change in rscu values:
ns_codon_counts <- na.omit(codon_counts_df[!(codon_counts_df$aa_major ==
                                              codon_counts_df$aa_minor) &
                                            !(codon_counts_df$major_codon ==
                                                codon_counts_df$minor_codon),])


### Contingency tables summing across codon pairs

# codons:
OR_codons <- ddply(ns_codon_counts, c("species", "codon_pair", "aa3_pair"),
```

```
                   function(x) {
  sum.cross_p <- sum(x$cross_p, na.rm = TRUE)
  sum.cross_u <- sum(x$cross_u, na.rm = TRUE)
  OR1 <- sum(x$cross_p, na.rm = TRUE)/sum(x$cross_u, na.rm = TRUE)
  OR2 <- sum(x$cross_p2, na.rm = TRUE)/sum(x$cross_u2, na.rm = TRUE)
  data.frame(sum.cross_p, sum.cross_u, OR1, OR2)
})

# log(OR):
OR_codons$log.OR1 <- log(OR_codons$OR1)
OR_codons$log.OR2 <- log(OR_codons$OR2)

# some codon pairs only have polymorphisms in one of the directions,
# so will remove these from analysis (Arg<>Pro)
OR_codons$log.OR1[is.infinite(OR_codons$log.OR1) == TRUE] <- NA
OR_codons$log.OR2[is.infinite(OR_codons$log.OR2) == TRUE] <- NA
OR_codons <- na.omit(OR_codons)

# adding log_rscu values for each codon pair
codon_rscu <- unique(subset(single_codon_pair, select = c("species", "codon_pair", "log_rscu")))
OR_codons <- merge(OR_codons, codon_rscu, by = c("species", "codon_pair"))
write.table(OR_codons, file = "~/Dropbox/CUB_supplementary_data/tables/OR_tbl_codons.csv",
            sep = "\t", col.names = T, row.names = F, quote = F)
```

Distribution of log(Y) across codons in each species.

```
# distribution of log(Y) (OR1)
OR_codons$species <- factor(OR_codons$species, levels = c("ecoli", "spneumoniae"))
levels(OR_codons$species) <- c(expression(italic("E. coli")),
                               expression(italic("S. pneumoniae")))

p.dist.OR1.codon <- ggplot(OR_codons, aes(x=log.OR1)) +
  geom_histogram(aes(y=..density..),
                 binwidth=.2,
                 colour="black", fill="white") +
  xlab(expression(paste("log(", italic("Y"), ")", sep = ""))) +
  ylab("Frequency") +
  geom_vline(data = ddply(OR_codons, "species", summarise, avg = mean(log.OR1)),
             aes(xintercept=avg),linetype="dashed",
             color = "azure4", size=0.5) +
  facet_grid(~species, scales = "free", labeller = label_parsed) +
  geom_text(data = ddply(OR_codons, "species", summarise, avg = mean(log.OR1)),
            aes(x = -3, y = 0.5, label = paste("mean(log(Y)) = ",
                                               round(avg, digits = 3), sep = "")),
            hjust = 0,family = "Times", size = 4, color = "azure4") +
  theme_bw() +
  theme(text = element_text(family = "Times", size = 14),
        axis.text = element_text(family = "Times", size = 14),
        strip.text.x = element_text(family = "Times", face = "bold", size = 14),
        strip.background.x = element_rect(fill = "gray90", linetype = "blank"))
p.dist.OR1.codon
```

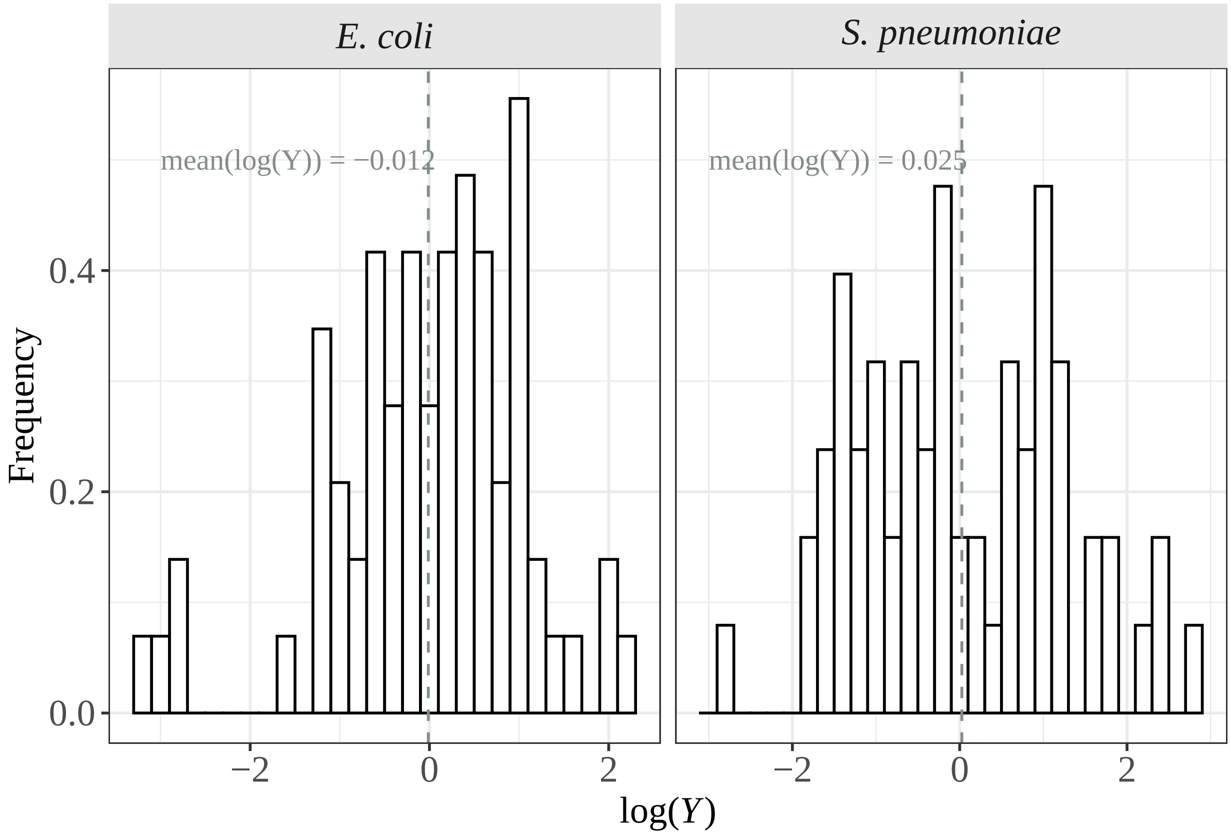


```
# checking if the mean is different form 0:
p_logY <- ddply(OR_codons, "species", function(x) {
  t_test <- t.test(x$log.OR1)
  mean <- t_test$estimate
  p_value <- t_test$p.value
  data.frame(mean, p_value)
})
kable(p_logY)
```

| species | mean | p_value |
|---|---|---|
| italic("E. coli") | -0.0124935 | 0.9235116 |
| italic("S. pneumoniae") | 0.0254538 | 0.8707573 |

```
# distribution of log(Y) (OR2)
p.dist.OR2.codon <- ggplot(OR_codons, aes(x=log.OR2)) +
  geom_histogram(aes(y=..density..),
               binwidth=.2,
               colour="black", fill="white") +
  xlab(expression(paste("log(", italic("Y"), ")", sep = ""))) +
  ylab("Frequency") +
  geom_vline(data = ddply(OR_codons, "species", summarise, avg = mean(log.OR2)),
           aes(xintercept=avg),linetype="dashed",
           color = "azure4", size=0.5) +
  facet_grid(~species, scales = "free", labeller = label_parsed) +
  geom_text(data = ddply(OR_codons, "species", summarise, avg = mean(log.OR2)),
          aes(x = -3, y = 0.8, label = paste("mean(log(Y)) = ",
```

```
                                                    round(avg, digits = 3), sep = "")),
             hjust = 0,family = "Times", size = 4, color = "azure4") +
  theme_bw() +
  theme(text = element_text(family = "Times", size = 14),
        axis.text = element_text(family = "Times", size = 14),
        strip.text.x = element_text(family = "Times", face = "bold", size = 14),
        strip.background.x = element_rect(fill = "gray90", linetype = "blank"))
p.dist.OR2.codon
```

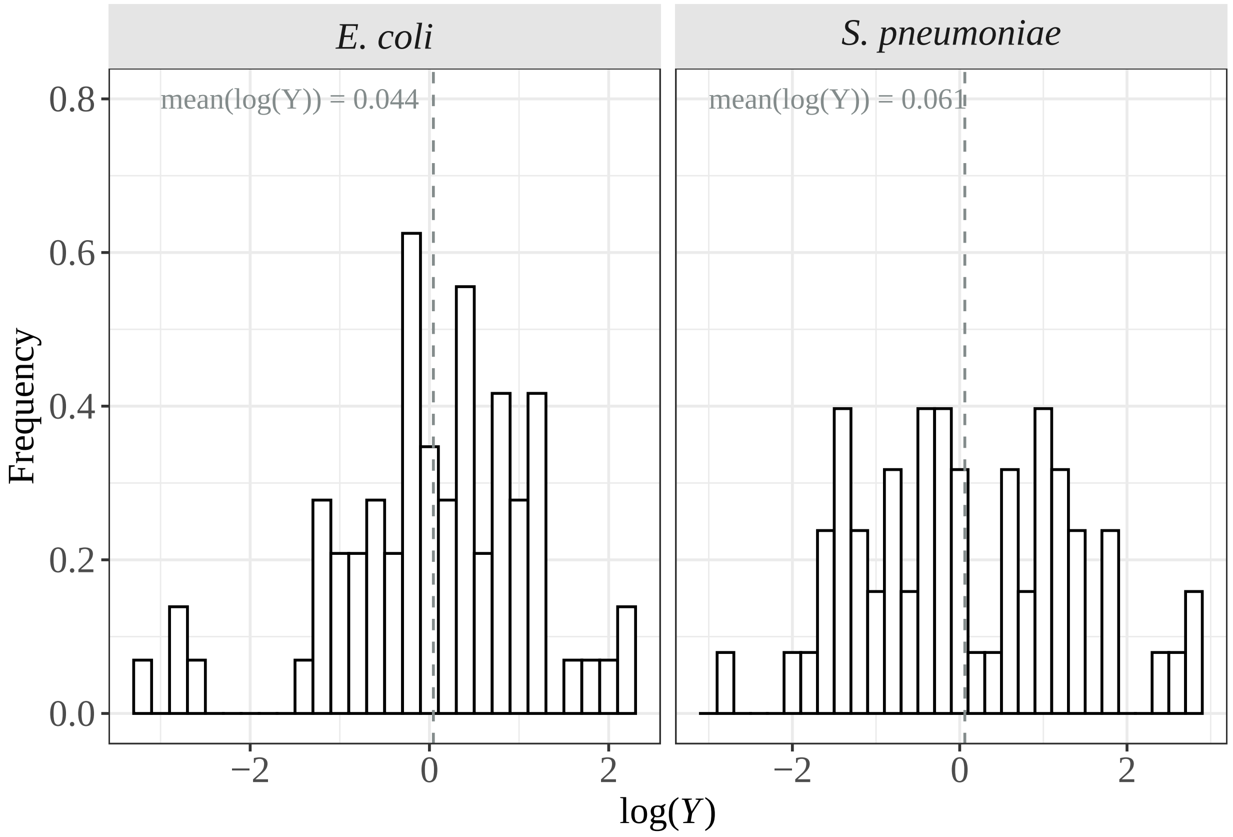


```
# checking if the mean is different form 0:
p_logY2 <- ddply(OR_codons, "species", function(x) {
  t_test <- t.test(x$log.OR2)
  mean <- t_test$estimate
  p_value <- t_test$p.value
  data.frame(mean, p_value)
})
kable(p_logY2)
```

| species | mean | p_value |
|---|---|---|
| italic("E. coli") | 0.043856 | 0.7367237 |
| italic("S. pneumoniae") | 0.061193 | 0.7042350 |

```
### NOTE: as results are very similar between log.OR1 and log.OR2, we will use log.OR1
```

Relationship between log(Y) and log(RSCUp/RSCUu).

```r
# log(Y) ~ log(rscu_p/rscu_u)
p.RSCU.OR1 <- ggplot(OR_codons, aes(log_rscu, log.OR1, label = codon_pair)) +
    geom_point(size = 0.6) +
    geom_smooth(method = "glm", formula = y~x, se = F, color = "black", size = 1) +
    xlab(expression(paste("log(", italic(RSCU[p]/RSCU[u]), ")", sep = ""))) +
    ylab(expression(paste("log(", italic("Y"), ")", sep = ""))) +
    stat_cor(label.x = 2, label.y.npc = "bottom", method = "pearson", size = 3.5) +
    facet_grid(~species, scales = "free", labeller = label_parsed) +
    theme_bw() +
    theme(text = element_text(family = "Times", size = 14),
        strip.text.x = element_text(family = "Times", size = 14),
        strip.text.y = element_text(family = "Times", face = "bold", size = 14),
        strip.background.x = element_rect(fill = "gray90", linetype = "blank"),
        strip.background.y = element_rect(fill = "gray90", linetype = "blank"))
p.RSCU.OR1
```

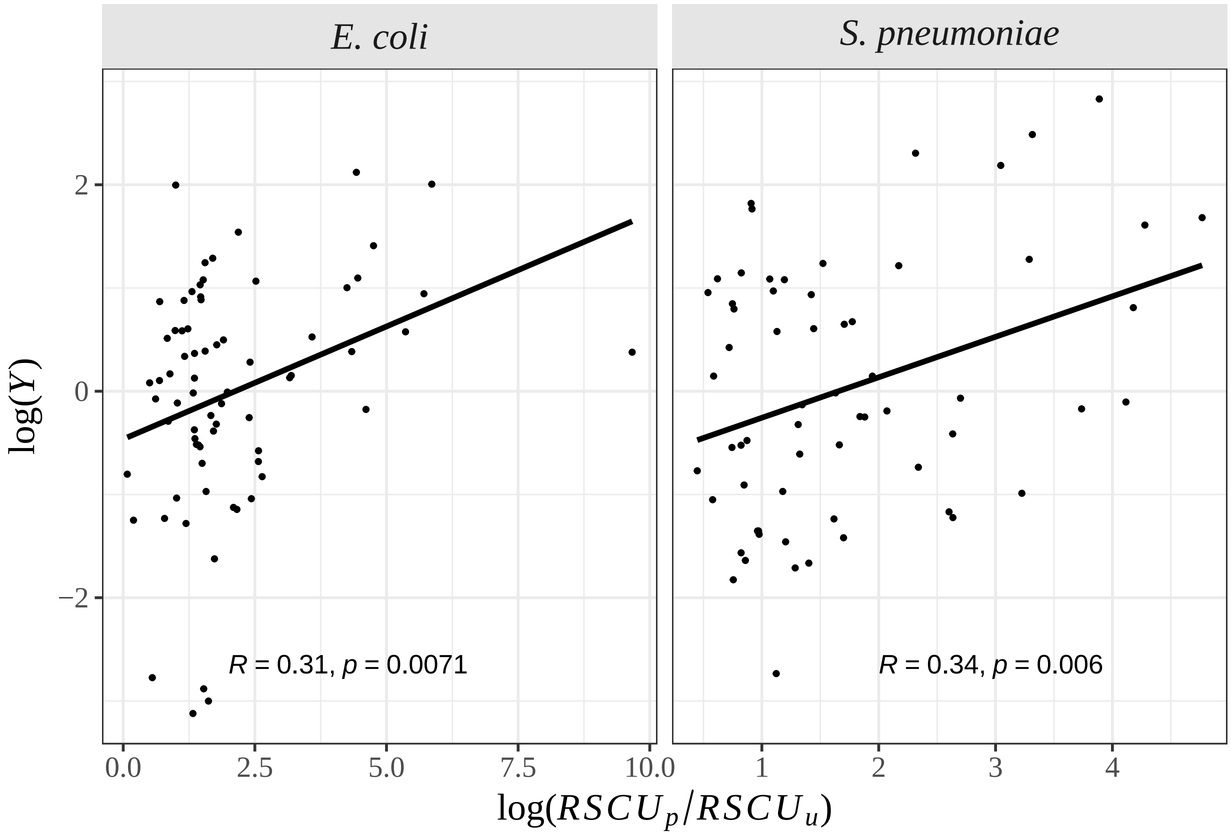


log(Y) aalysis at the gene level.

```r
### Contingency tables summing across genes

# gene:
OR_gene <- ddply(ns_codon_counts, c("species", "coID"), function(x) {
  sum.cross_p <- sum(x$cross_p, na.rm = TRUE)
  sum.cross_u <- sum(x$cross_u, na.rm = TRUE)
  OR1 <- sum(x$cross_p, na.rm = TRUE)/sum(x$cross_u, na.rm = TRUE)
  OR2 <- sum(x$cross_p2, na.rm = TRUE)/sum(x$cross_u2, na.rm = TRUE)
```

```
  data.frame(sum.cross_p, sum.cross_u, OR1, OR2)
})

# log(OR):
OR_gene$log.OR1 <- log(OR_gene$OR1)
OR_gene$log.OR2 <- log(OR_gene$OR2)

# some codon pairs only have polymorphisms in one of the directions,
# so will remove these from analysis
OR_gene$log.OR1[is.infinite(OR_gene$log.OR1) == TRUE] <- NA
OR_gene$log.OR2[is.infinite(OR_gene$log.OR2) == TRUE] <- NA
OR_gene <- na.omit(OR_gene)

# adding cai and gene expression data:
cai_exp <- read.table(file = "~/Dropbox/CUB_supplementary_data/tables/gene_cai_exp.csv",
                      sep = "\t", header = T)
OR_gene <- merge(OR_gene, cai_exp, by = c("coID", "species"))
write.table(OR_gene, file = "~/Dropbox/CUB_supplementary_data/tables/OR_tbl_gene.csv",
            sep = "\t", col.names = T, row.names = F, quote = F)


# distribution of log(Y):
OR_gene$species <- factor(OR_gene$species, levels = c("ecoli", "spneumoniae"))
levels(OR_gene$species) <- c(expression(italic("E. coli")),
                             expression(italic("S. pneumoniae")))

p.dist.OR1.gene <- ggplot(OR_gene, aes(x=log.OR1)) +
  geom_histogram(aes(y=..density..),
                 binwidth=.2,
                 colour="black", fill="white") +
  xlab(expression(paste("log(", italic("Y"), ")", sep = ""))) +
  ylab("Frequency") +
  geom_vline(data = ddply(OR_gene, "species", summarise, avg = mean(log.OR1)),
             aes(xintercept=avg),linetype="dashed",
             color = "azure4", size=0.5) +
  facet_grid(~species, scales = "free", labeller = label_parsed) +
  geom_text(data = ddply(OR_gene, "species", summarise, avg = mean(log.OR1)),
            aes(x = -4, y = 0.5, label = paste("mean(log(Y)) = ",
                                                round(avg, digits = 3), sep = "")),
            hjust = 0,family = "Times", size = 4, color = "azure4") +
  theme_bw() +
  theme(text = element_text(family = "Times", size = 14),
        axis.text = element_text(family = "Times", size = 14),
        strip.text.x = element_text(family = "Times", face = "bold", size = 14),
        strip.background.x = element_rect(fill = "gray90", linetype = "blank"))
p.dist.OR1.gene
```

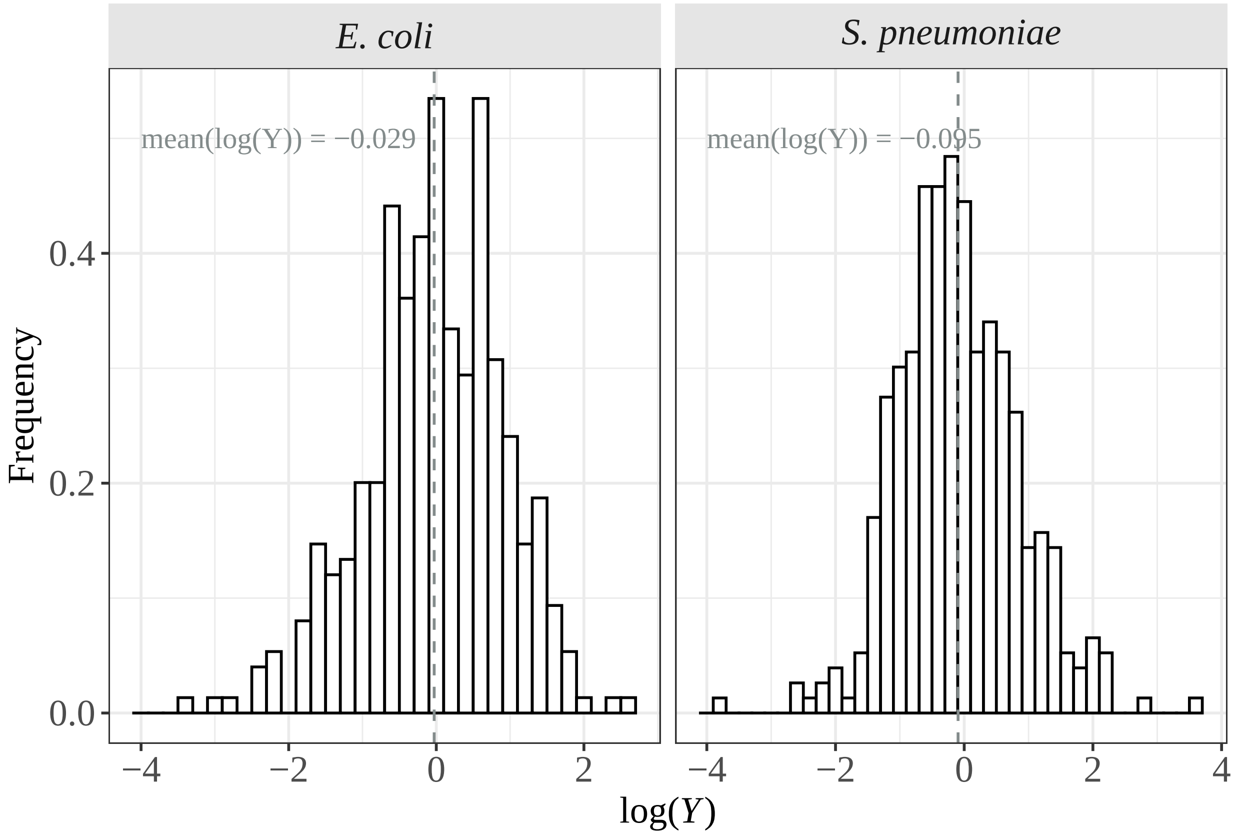


```
# checking if the mean is different form 0:
p_logY <- ddply(OR_gene, "species", function(x) {
  t_test <- t.test(x$log.OR1)
  mean <- t_test$estimate
  p_value <- t_test$p.value
  data.frame(mean, p_value)
})
kable(p_logY)
```

| species | mean | p_value |
|---|---|---|
| italic("E. coli") | -0.0287375 | 0.5556791 |
| italic("S. pneumoniae") | -0.0952318 | 0.0474858 |

```
# log(Y) ~ CAI:
p.cai.OR1 <- ggplot(OR_gene, aes(cai_major, log.OR1, label = coID)) +
  geom_point(size = 0.6) +
  geom_smooth(method = "glm", formula = y~x, se = F, color = "black", size = 1) +
  xlab("CAI") +
  ylab(expression(paste("log(", italic("Y"), ")", sep = ""))) +
  stat_cor(label.x.npc = "left", label.y.npc = "top", method = "pearson", size = 3.5) +
  facet_grid(~species, scales = "free", labeller = label_parsed) +
  theme_bw() +
  theme(text = element_text(family = "Times", size = 14),
        axis.text = element_text(family = "Times", size = 14),
        strip.text.x = element_text(family = "Times", face = "bold", size = 14),
```

```
        strip.background.x = element_rect(fill = "gray90", linetype = "blank"))
p.cai.OR1
```

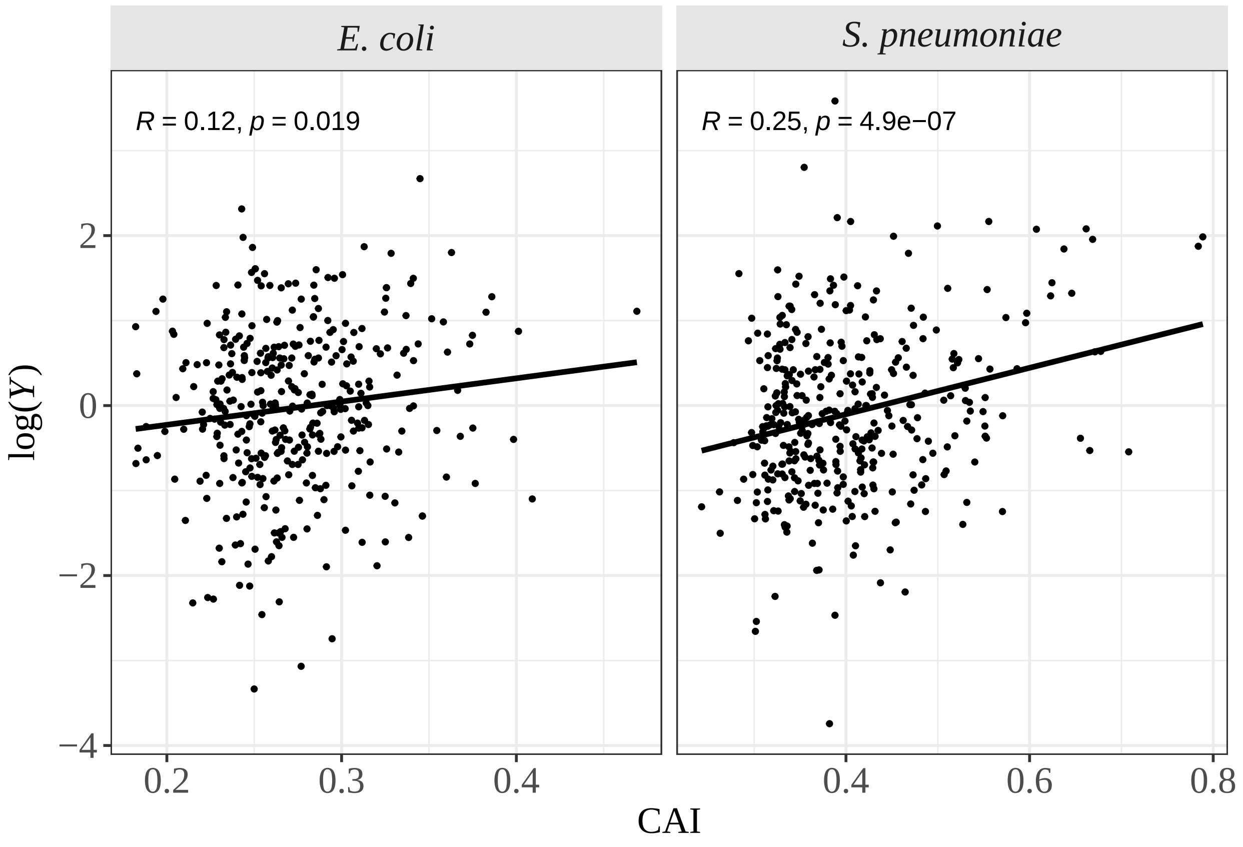


```
# log(Y) ~ gene expression
p.exp.OR1 <- ggplot(OR_gene, aes(mean_exp, log.OR1, label = coID)) +
  geom_point(size = 0.6) +
  geom_smooth(method = "glm", formula = y~x, se = F, color = "black", size = 1) +
  xlab("log(gene expression)") +
  ylab(expression(paste("log(", italic("Y"), ")", sep = ""))) +
  stat_cor(label.x.npc = "left", label.y.npc = "top", method = "pearson", size = 3.5) +
  facet_grid(~species, scales = "free", labeller = label_parsed) +
  theme_bw() +
  theme(text = element_text(family = "Times", size = 14),
        axis.text = element_text(family = "Times", size = 14),
        strip.text.x = element_text(family = "Times", face = "bold", size = 14),
        strip.background.x = element_rect(fill = "gray90", linetype = "blank"))
p.exp.OR1
```

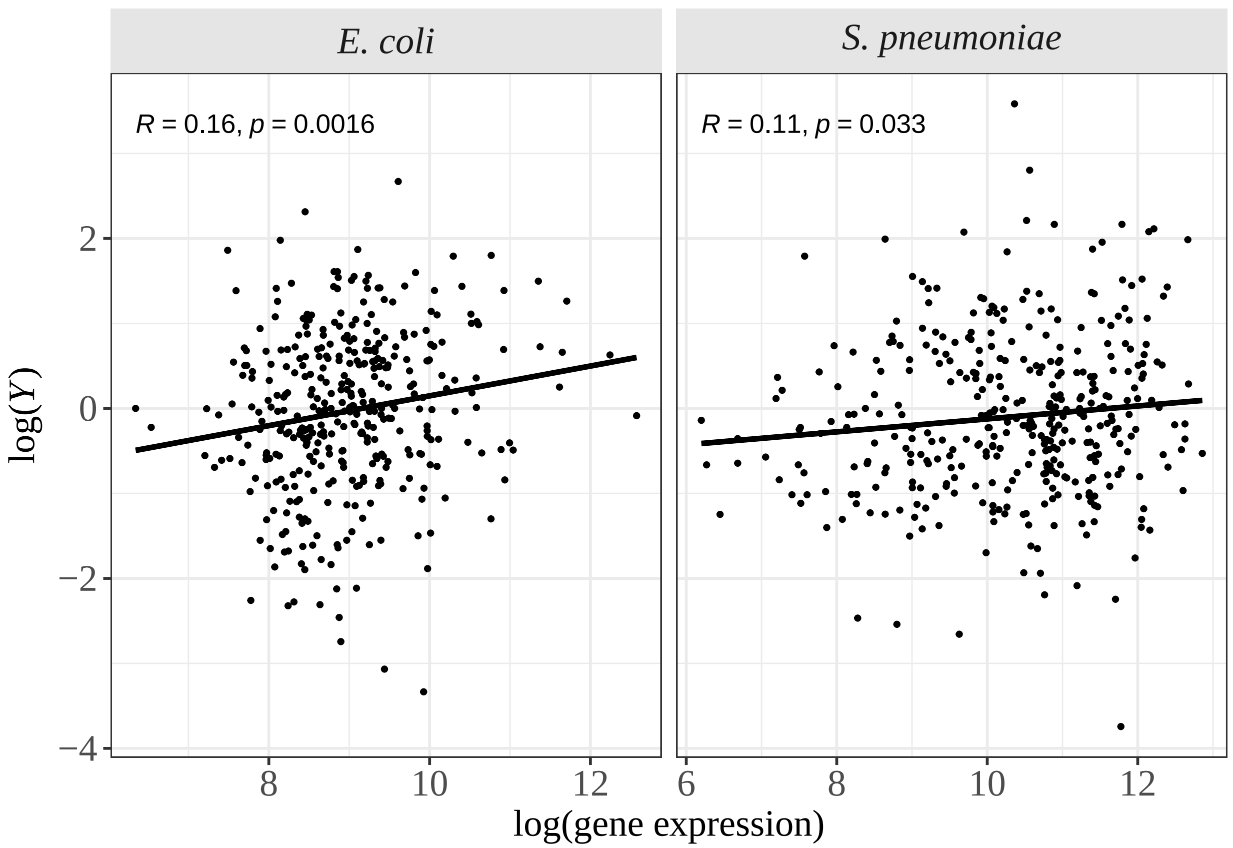
E. coli
S. pneumoniae
R = 0.16, p = 0.0016
R = 0.11, p = 0.033
log(Y)
2
0
−2
−4
8
10
12
6
8
10
12
log(gene expression)
