## Supplementary material for "The silent impact: codon usage bias and protein evolution in bacteria": File S2

### Divergence analysis

af_moutinho

2022-07-04

This RMarkdown contains all analyses performed at the divergence level. The first chunk of the script reproduces the estimation of Z in *E. coli* and *S. pneumoniae* using the RSCU values of highly expressed genes.

```
# Libraries
library(plyr)
library(dplyr)
library(data.table)
library(ggplot2)
library(ggpubr)
library(kableExtra)
#

# table contanining all possible codon pairs for E. coli and S. pneumoniae
# using the RSCU value for highly expressed genes
codon_subs_high <- fread(file = "~/Dropbox/CUB_supplementary_data/tables/codon_subs_rscu_high.csv",
                         sep = "\t", header = T)

# keeping only codon pairs with 1 nucleotide difference and only those involving
# non-synonymous substitutions:
ns_codon_subs_high <- subset(codon_subs_high, nt.dif == 1 & !(aa1 == aa2))

# estimating the weighted averages:
# NOTE: adding 1e-04 to also consider codons where RSCU is 0
ns_codon_subs_high$weight1 <- with(ns_codon_subs_high,
                                   counts1*(log((rscu2 + 1e-04)/(rscu1 + 1e-04))))

# summing across codon pairs for each amino acid pair:
sum_weights_high <- ddply(ns_codon_subs_high, c("aa_pair", "species"),
                          function(x) {
  Z = sum(x$weight1, na.rm = TRUE)/sum(x$counts1, na.rm = TRUE)
  data.frame(Z)
})

# putting the species name in italic:
sum_weights_plot <- sum_weights_high # doing another table as the species names will be in italic
sum_weights_plot$species <- factor(sum_weights_plot$species,
                                   levels = c("ecoli", "spneumoniae"))
levels(sum_weights_plot$species) <- c(expression(italic("E. coli")),
                                      expression(italic("S. pneumoniae")))

# plotting the distribution of Z values:
```

```
p.dist.Z_high <- ggplot(sum_weights_plot, aes(x=Z)) +
  geom_histogram(aes(y=..density..),
                 binwidth=.2,
                 colour="black", fill="white") +
  xlab(expression(italic("Z"))) +
  ylab("Frequency") +
  geom_vline(data = ddply(sum_weights_plot, c("species"), summarise, avg = mean(Z)),
             aes(xintercept=avg),linetype="dashed",
             color = "azure4", size=0.5) +
  facet_grid(.~species, scales = "free", labeller = label_parsed) +
  geom_text(data = ddply(sum_weights_plot, c("species"), summarise, avg = mean(Z)),
            aes(x = -5, y = 0.9, label = paste("mean(Z) = ", round(avg, digits = 3), sep = "")),
            hjust = 0,family = "Times", size = 4, color = "azure4") +
  theme_bw() +
  theme(text = element_text(family = "Times", size = 14),
        axis.text = element_text(family = "Times", size = 14),
        strip.text.x = element_text(family = "Times", size = 14),
        strip.text.y = element_text(family = "Times", face = "bold", size = 14),
        strip.background.x = element_rect(fill = "gray90", linetype = "blank"),
        strip.background.y = element_rect(fill = "gray90", linetype = "blank"))

p.dist.Z_high
```

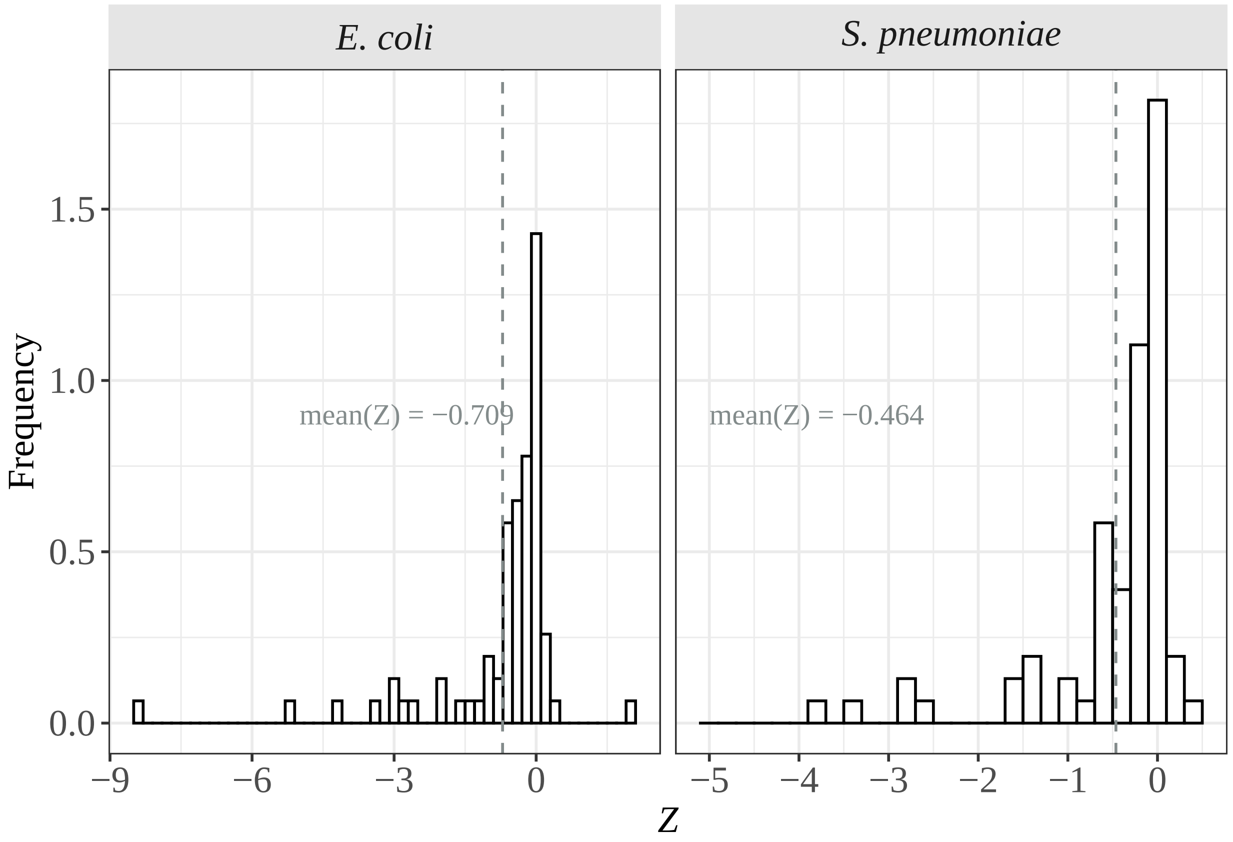


The next chunk of the script contains the analysis of the relationship between $Z$ and the divergence between amino aicds ($dij$)

```r
# table with divergence estimates for each amino acid pair in each species
dxy_df <- read.table(file = "~/Dropbox/CUB_supplementary_data/tables/dxy_df.csv",
                     sep = "\t", header = T)

# cobining the two tables
sum_weights_high <- merge(sum_weights_high, dxy_df, by = c("species", "aa_pair"))

# species names in italic
sum_weights_high$species <- factor(sum_weights_high$species,
                                   levels = c("ecoli", "spneumoniae"))
levels(sum_weights_high$species) <- c(expression(italic("E. coli")),
                                     expression(italic("S. pneumoniae")))

p.Z.dxy <- ggplot(sum_weights_high, aes(Z, dxy, label = aa_pair)) +
  geom_point(size = 0.6) +
  geom_smooth(method = "lm", formula = y~x, se = F, color = "black", size = 1) +
  xlab(expression(italic(Z))) +
  ylab(expression(italic(d[ij]))) +
  stat_cor(label.x.npc = "left", label.y.npc = "top", method = "pearson", size = 3.5) +
  facet_grid(.~species, scales = "free", labeller = label_parsed) +
  theme_bw() +
  theme(text = element_text(family = "Times", size = 14),
        strip.text.x = element_text(family = "Times", size = 14),
        strip.text.y = element_text(family = "Times", face = "bold", size = 14),
        strip.background.x = element_rect(fill = "gray90", linetype = "blank"),
        strip.background.y = element_rect(fill = "gray90", linetype = "blank"))
p.Z.dxy
```

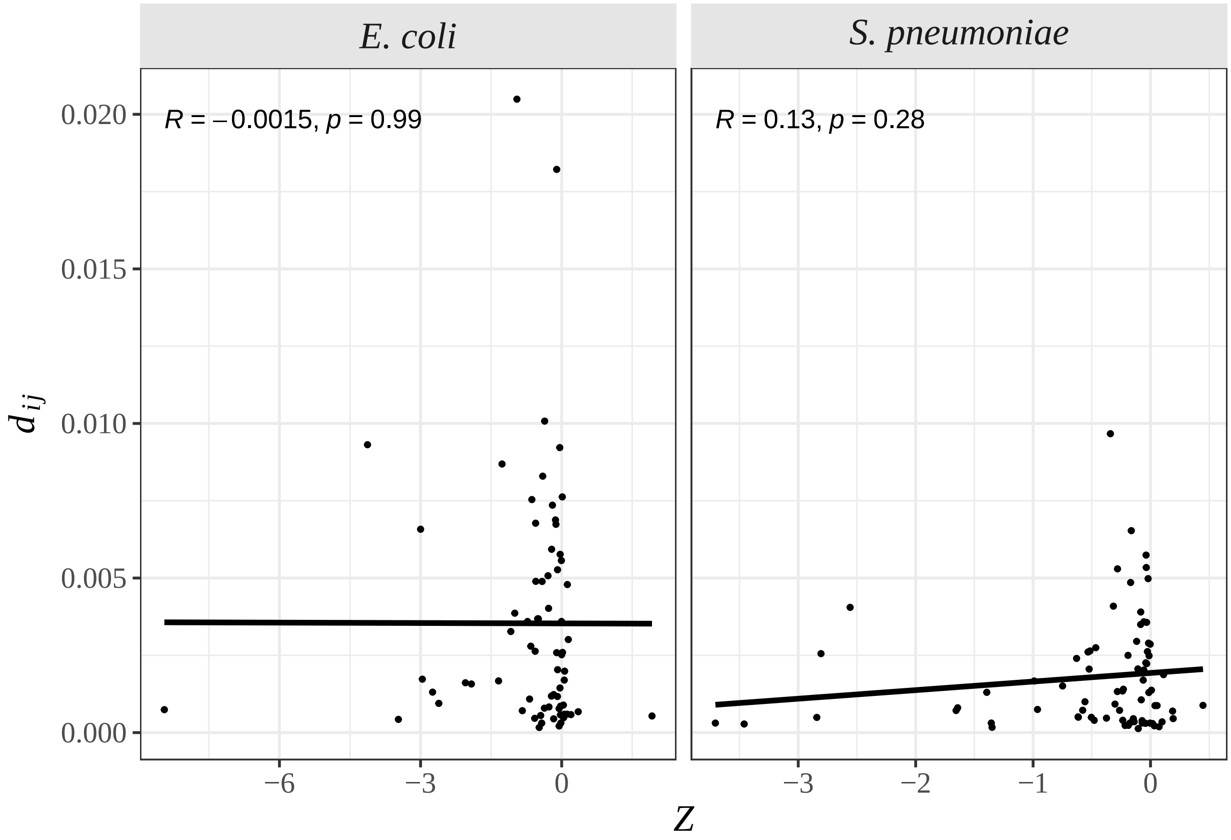


Analysis of Z for each category of CAI (Codon Adaptation Index)

```
# this analysis will performed using the RSCU values estimated from each CAI category

codon_subs_cai <-
  read.table(file = "~/Dropbox/CUB_supplementary_data/tables/codons_subs_dxy_cai_tbl.csv",
                              sep = "\t", header = T)

# summing all codons in each contig for each species
counts_codon_subs <- ddply(codon_subs_cai, c("species", "coID", "cai_cat", "codon1"),
                      function(x) {
  counts1 <- sum(unique(x$counts1_all))
  data.frame(counts1)
})

# summing all occurrences of each codon:
counts_codon_cai <- ddply(counts_codon_subs, c("species", "codon1", "cai_cat"), function(x) {
  counts1 <- sum(x$counts1)
  data.frame(counts1)
})

# taking only non-synonymous aa pairs seperated by 1 mutation
ns_codons <- subset(codon_subs_cai, nt.dif == 1 & !aa1 == aa2)

df_ns_codons <- unique(subset(ns_codons, select = c("cai_cat", "codon2", "codon1", "species",
                                                    "aa_pair", "rscu1", "rscu2")))
```

```r
# combing with the table containing all counts of each codon:
df2_ns_codons <- merge(df_ns_codons, counts_codon_cai, by = c("species", "cai_cat",
                                                              "codon1"))

# estimating the weighted averages:
df2_ns_codons$weight1 <- df2_ns_codons$counts1*(log((df2_ns_codons$rscu2 + 1e-04)/
                                                      (df2_ns_codons$rscu1 + 1e-04)))

sum_weights_cai <- ddply(df2_ns_codons, c("species", 'cai_cat', "aa_pair"), function(x) {
  Z = sum(x$weight1, na.rm = TRUE)/sum(unique(x$counts1), na.rm = TRUE)
  data.frame(Z)
})

# adding the dxy estimates for each amino acid pair:
sum_weights_cai <- merge(sum_weights_cai, dxy_df, by = c("aa_pair", "species"))

sum_weights_cai$cai_cat <- as.character(sum_weights_cai$cai_cat)

# species names in italic
sum_weights_cai$species <- factor(sum_weights_cai$species,
                                  levels = c("ecoli", "spneumoniae"))
levels(sum_weights_cai$species) <- c(expression(italic("E. coli")),
                                     expression(italic("S. pneumoniae")))

p.Z.cai.dxy <- ggplot(sum_weights_cai,
                      aes(Z, dxy, label = aa_pair, colour = cai_cat)) +
  geom_point(size = 0.5) +
  geom_smooth(method = "glm", formula = y~x, se = FALSE) +
  xlab(expression(italic(Z))) +
  ylab(expression(italic(d[ij]))) +
  #geom_quantile(formula=y~x) +
  guides(col=guide_legend(title="CAI", label = T)) +
  facet_wrap(~species, scales = "free", labeller = label_parsed) +
  scale_color_viridis_d(label = c(expression(1^st), expression(2^nd),
                                  expression(3^rd), expression(4^th))) +
  theme_bw() +
  theme_bw() +
  theme(text = element_text(family = "Times", size = 14),
        strip.text.x = element_text(family = "Times", size = 14),
        strip.text.y = element_text(family = "Times", face = "bold", size = 14),
        strip.background.x = element_rect(fill = "gray90", linetype = "blank"),
        strip.background.y = element_rect(fill = "gray90", linetype = "blank"))
p.Z.cai.dxy
```

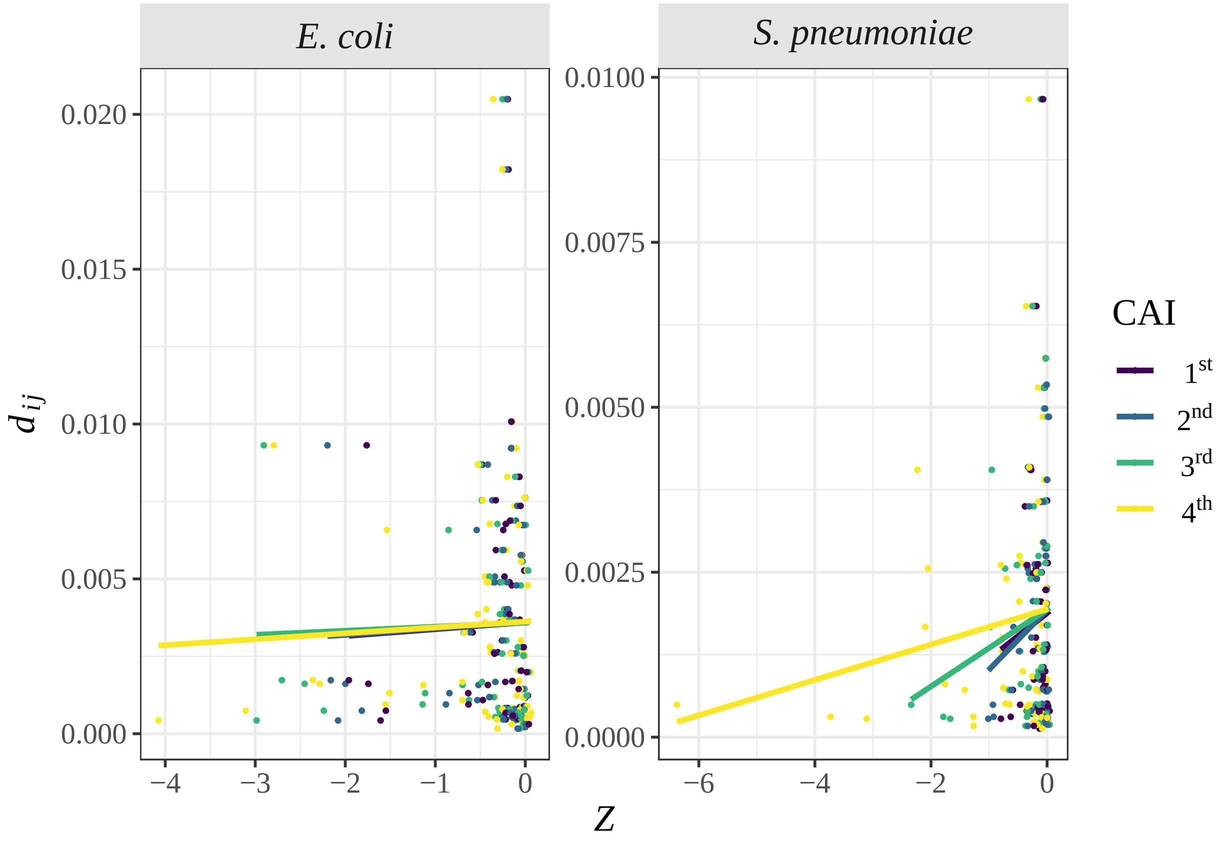


```r
# correlation between Z and dij for each CAI category:
p_Z_cai <- ddply(sum_weights_cai, c("species", "cai_cat"), function(x) {
  cor_df <- cor.test(x$Z, x$dxy, method = "pearson", exact = F)
  estimate <- cor_df$estimate
  p_value <- cor_df$p.value
  data.frame(estimate, p_value)
})
kable(x = p_Z_cai)
```

| species | cai_cat | estimate | p_value |
|---|---|---|---|
| italic("E. coli") | 1 | 0.0247349 | 0.8377660 |
| italic("E. coli") | 2 | 0.0275877 | 0.8193547 |
| italic("E. coli") | 3 | 0.0230262 | 0.8488364 |
| italic("E. coli") | 4 | 0.0390224 | 0.7466227 |
| italic("S. pneumoniae") | 1 | 0.0618212 | 0.5932602 |
| italic("S. pneumoniae") | 2 | 0.1086929 | 0.3467212 |
| italic("S. pneumoniae") | 3 | 0.1320780 | 0.2521894 |
| italic("S. pneumoniae") | 4 | 0.1452741 | 0.2074443 |

The last chunk of the script reproduces the estimation of Z in E. coli and S. pneumoniae using the RSCU values estimated using the whole genome:

```r
# table containining all possible codon pairs for E. coli and S. pneumoniae
# using the RSCU value estimated from the full genome
codon_subs_all <- fread(file = "~/Dropbox/CUB_supplementary_data/tables/codon_subs_rscu_all.csv",
                        sep = "\t", header = T)
```

```r
# keeping only codon pairs with 1 nucleotide difference and only those
# involving non-synonymous substitutions:
ns_codon_subs_all <- subset(codon_subs_all, nt.dif == 1 & !(aa1 == aa2))

# estimating the weighted averages:
# NOTE: adding 1e-04 to also consider codons where RSCU is 0
ns_codon_subs_all$weight1 <- with(ns_codon_subs_all, counts1_all*(log((rscu2_all + 1e-04)/
                                                                    (rscu1_all + 1e-04))))

# summing across codon pairs for each amino acid pair:
sum_weights_all <- ddply(ns_codon_subs_all, c("aa_pair", "species"), function(x) {
  Z = sum(x$weight1, na.rm = TRUE)/sum(x$counts1, na.rm = TRUE)
  data.frame(Z)
})

# adding dxy values:
sum_weights_all <- merge(sum_weights_all, dxy_df, by = c("species", "aa_pair"))

# putting the species name in italic:
sum_weights_all$species <- factor(sum_weights_all$species,
                                  levels = c("ecoli", "spneumoniae"))
levels(sum_weights_all$species) <- c(expression(italic("E. coli")),
                                     expression(italic("S. pneumoniae")))

# plotting the distribution of Z values:
p.dist.Z_all <- ggplot(sum_weights_all, aes(x=Z)) +
  geom_histogram(aes(y=..density..),
                 binwidth=.2,
                 colour="black", fill="white") +
  xlab(expression(italic("Z"))) +
  ylab("Frequency") +
  geom_vline(data = ddply(sum_weights_all, c("species"),
                          summarise, avg = mean(Z)),
             aes(xintercept=avg),linetype="dashed",
             color = "azure4", size=0.5) +
  facet_grid(.~species, scales = "free", labeller = label_parsed) +
  geom_text(data = ddply(sum_weights_all, c("species"),
                         summarise, avg = mean(Z)),
            aes(x = -5, y = 0.9, label = paste("mean(Z) = ", round(avg, digits = 3),
                                               sep = "")),
            hjust = 0,family = "Times", size = 4, color = "azure4") +
  theme_bw() +
  theme(text = element_text(family = "Times", size = 14),
        axis.text = element_text(family = "Times", size = 14),
        strip.text.x = element_text(family = "Times", size = 14),
        strip.text.y = element_text(family = "Times", face = "bold", size = 14),
        strip.background.x = element_rect(fill = "gray90", linetype = "blank"),
        strip.background.y = element_rect(fill = "gray90", linetype = "blank"))

p.dist.Z_all
```

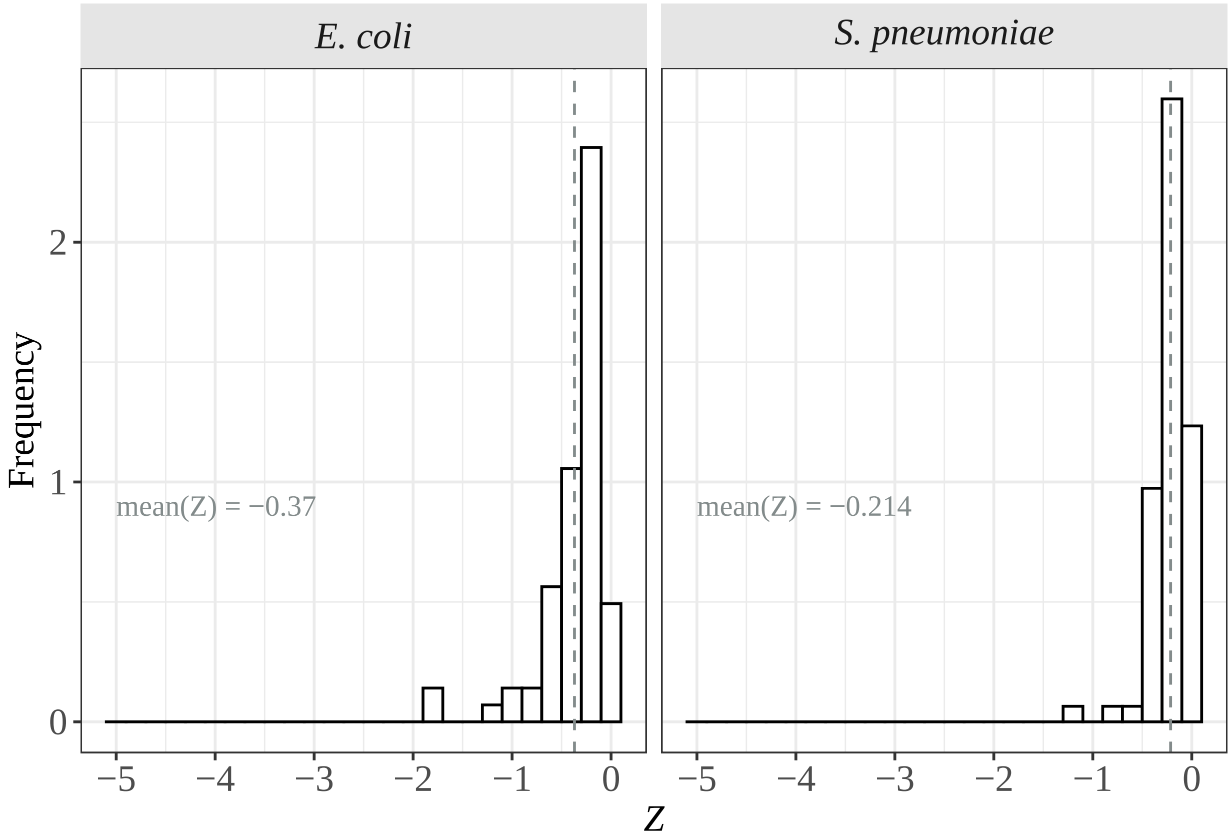


The next chunk of the script contains the analysis of the relationship between Z and the divergence between amino aicds (dij) using RSCU values for the full genome.

```
# Z ~ dxy
p.Z.dxy_all <- ggplot(sum_weights_all, aes(Z, dxy, label = aa_pair)) +
  geom_point(size = 0.6) +
  geom_smooth(method = "lm", formula = y~x, se = F, color = "black", size = 1) +
  xlab(expression(italic(Z))) +
  ylab(expression(italic(d[ij]))) +
  stat_cor(label.x.npc = "left", label.y.npc = "top", method = "pearson", size = 3.5) +
  facet_grid(.~species, scales = "free", labeller = label_parsed) +
  theme_bw() +
  theme(text = element_text(family = "Times", size = 14),
        strip.text.x = element_text(family = "Times", size = 14),
        strip.text.y = element_text(family = "Times", face = "bold", size = 14),
        strip.background.x = element_rect(fill = "gray90", linetype = "blank"),
        strip.background.y = element_rect(fill = "gray90", linetype = "blank"))
p.Z.dxy_all
```

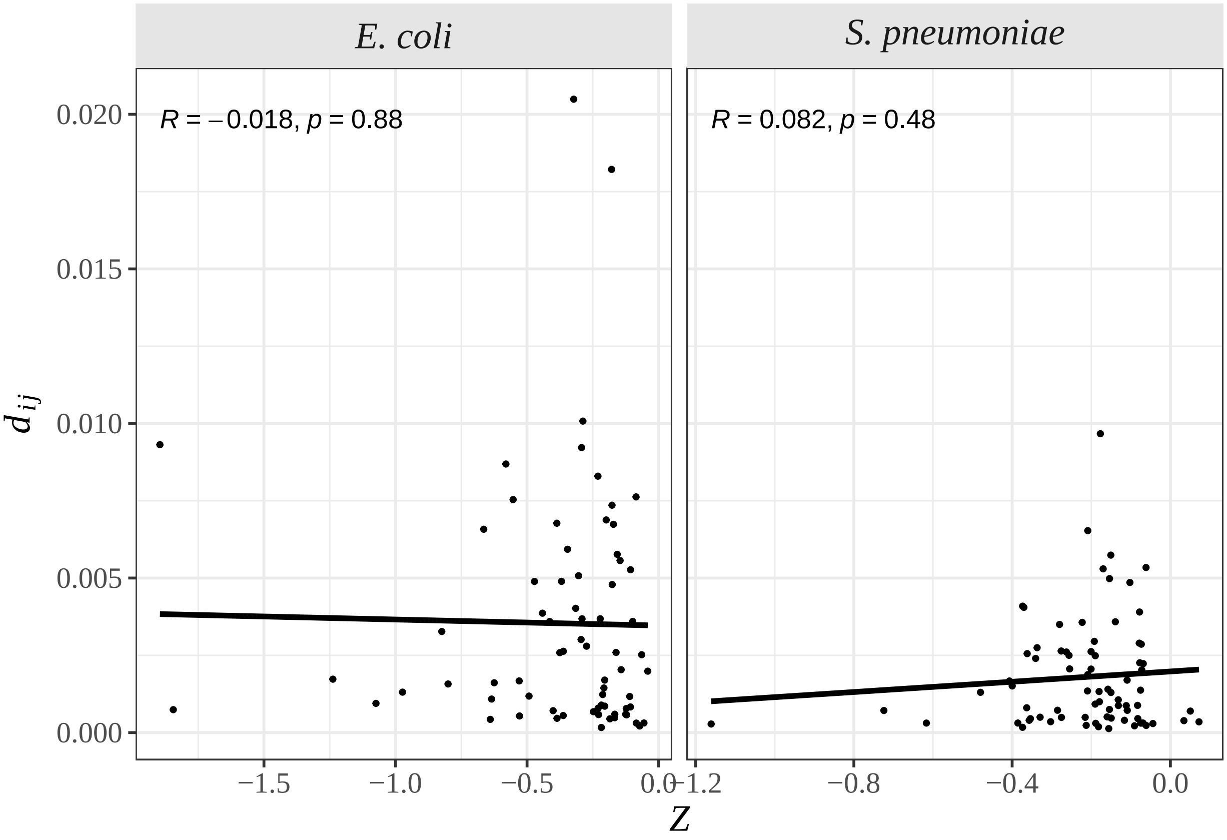
E. coli
S. pneumoniae
R = −0.018, p = 0.88
R = 0.082, p = 0.48
0.020
0.015
0.010
0.005
0.000
−1.5
−1.0
−0.5
0.0
−1.2
−0.8
−0.4
0.0
d_ij
Z
