## Supplementary material for "The silent impact: codon usage bias and protein evolution in bacteria": File S3

### Biased gene conversion and mutation bias analyses

af_moutinho

2022-07-05

This RMarkdown contains all analyses performed to assess the impact of biased gene conversion and mutation bias on selection acting on synonymous codon usage. This analysis was perfomed both at the polymorphism and substitution levels, where we subset the data into the following categories:

- **no-bias**: codon pairs not affected by biased gene conversion (A<>T and G<>C)
- **pAT**: codon pairs for which A or T are the preferred allele (i.e., GC are the unpreferred and AT the preferred alleles)
- **pGC**: codon pairs for which G or C are the preferred allele (i.e., AT are the unpreferred and GC the preferred alleles)

The following script contains the analyses performed at the polymorphism level.

```
# Libraries
library(data.table)
library(dplyr)
library(plyr)
library(ggplot2)
library(stringi)
library(stringr)
library(ggpubr)
library(kableExtra)
#

# getting the preferred/unpreferred codons from the polymorphism table:
codons_poly <- fread(file = "~/Dropbox/CUB_supplementary_data/tables/polymorphism_codons_tbl.csv",
                     sep = "\t", header = T)

# keeping only monomorphic/biallelic sites and codon pairs separated by 1 nt difference:
codons_poly2 <- codons_poly[codons_poly$nt.dif == 1 & codons_poly$NbAlleles == 2,]

# adding a column defining the preferred allele (p_allele) and unpreferred allele (u_allele)
codons_poly2$p_codon <- ifelse(codons_poly2$minor_codon == "p",
                               codons_poly2$MinorAllele, codons_poly2$MajorAllele)
codons_poly2$u_codon <- ifelse(codons_poly2$minor_codon == "np",
                               codons_poly2$MinorAllele, codons_poly2$MajorAllele)

# function to add the allele that differs in the p and u codon:
nt_allele.fun <- function(tbl) {
  for (i in 1:nrow(tbl)) {
    aa1 <- strsplit(tbl[i, "p_codon"], split = "")
```

```r
    aa2 <- strsplit(tbl[i, "u_codon"], split = "")
    comp.aa <- stri_compare(aa1[[1]], aa2[[1]])
    j <- which(!comp.aa == 0)
    tbl[i, "p_allele"] <- c(aa1[[1]][j], aa2[[1]][j])[1]
    tbl[i, "u_allele"] <- c(aa1[[1]][j], aa2[[1]][j])[2]
  }
  return(tbl)
}

codons_poly3 <- ddply(codons_poly2, "codon_pair", nt_allele.fun)

# taking the columns of interest:
codon_type <- unique(subset(codons_poly3, select = c("codon_pair", "p_codon", "u_codon",
                                                     "p_allele", "u_allele", "species")))

# getting the table with the log(Y) data for each codon site:
OR_codons <- fread(file = "~/Dropbox/CUB_supplementary_data/tables/OR_tbl_codons.csv",
                   sep = "\t", header = T)

# adding the mutation to each codon pair:
OR_codons$codon1 <- str_split_fixed(OR_codons$codon_pair, "-", 2)[,1]
OR_codons$codon2 <- str_split_fixed(OR_codons$codon_pair, "-", 2)[,2]

# combining the two tables:
OR_codons <- merge(OR_codons, codon_type, by = c("species", "codon_pair"))

# function that adds the mutation
nt_change.fun <- function(tbl) {
  for (i in 1:nrow(tbl)) {
    aa1 <- strsplit(tbl[i, "codon1"], split = "")
    aa2 <- strsplit(tbl[i, "codon2"], split = "")
    comp.aa <- stri_compare(aa1[[1]], aa2[[1]])
    j <- which(!comp.aa == 0)
    tbl[i, "mutation"] <- paste(sort(c(aa1[[1]][j], aa2[[1]][j]))[1],
                                sort(c(aa1[[1]][j], aa2[[1]][j]))[2], sep = "")
  }
  return(tbl)
}

OR_codons2 <-  ddply(OR_codons, "codon_pair", nt_change.fun)

# adding a column defining the mutation type:
OR_codons2$mutation_type[with(OR_codons2, (p_allele == "A" | p_allele == "T") &
                                (u_allele == "G" | u_allele == "C"))] <- "pAT"
OR_codons2$mutation_type[with(OR_codons2, (p_allele == "G" | p_allele == "C") &
                                (u_allele == "A" | u_allele == "T"))] <- "pGC"
OR_codons2$mutation_type[with(OR_codons2, mutation == "AT" | mutation == "CG")] <- "no-bias"
# savinf the new table with additional information
write.table(OR_codons2, file = "~/Dropbox/CUB_supplementary_data/tables/OR_tbl2_codons.csv",
            sep = "\t", col.names = T, row.names = F, quote = F)


### Distribution of log(Y) across different types of mutations
```

```
OR_codons2$species <- factor(OR_codons2$species, levels = c("ecoli", "spneumoniae"))
levels(OR_codons2$species) <- c(expression(italic("E. coli")),
                                expression(italic("S. pneumoniae")))

# logY ~ mutation
p.OR_mutation <- ggplot(OR_codons2, aes(x=log.OR1)) +
  geom_histogram(aes(y=..density..),
                 binwidth=.2,
                 colour="black", fill="white") +
  xlab(expression(paste("log(", italic("Y"), ")", sep = ""))) +
  ylab("Frequency") +
  geom_vline(data = ddply(OR_codons2, c("species", "mutation_type"), summarise,
                          avg = mean(log.OR1)),
             aes(xintercept=avg),linetype="dashed",
             color = "azure4", size=0.5) +
  facet_grid(species~mutation_type, scales = "free", labeller = label_parsed) +
  geom_text(data = ddply(OR_codons2, c("species", "mutation_type"),
                         summarise, avg = mean(log.OR1)),
            aes(x = -4, y = 1.15, label = paste("mean(log(Y)) = ",
                                                round(avg, digits = 3), sep = "")),
            hjust = 0,family = "Times", size = 4, color = "azure4") +
  theme_bw() +
  theme(text = element_text(family = "Times", size = 14),
        axis.text = element_text(family = "Times", size = 14),
        strip.text.x = element_text(family = "Times", size = 14),
        strip.text.y = element_text(family = "Times", face = "bold", size = 14),
        strip.background.x = element_rect(fill = "gray90", linetype = "blank"),
        strip.background.y = element_rect(fill = "gray90", linetype = "blank"))
p.OR_mutation
```

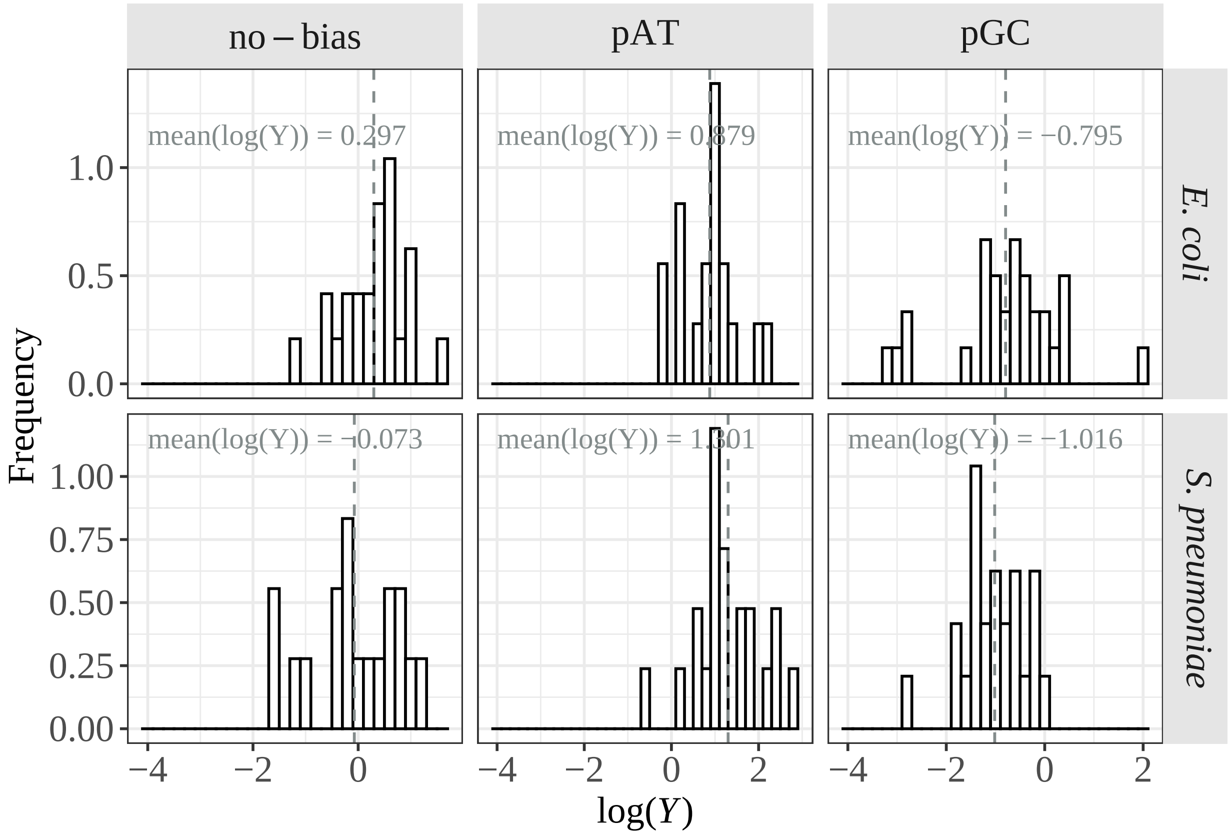


```
# checking if the mean is different form 0:
P_codon <- ddply(OR_codons2, c("species", "mutation_type"), function(x) {
  t_test <- t.test(x$log.OR1)
  mean <- t_test$estimate
  p_value <- t_test$p.value
  data.frame(mean, p_value)
})
kable(P_codon)
```

| species | mutation_type | mean | p_value |
|---|---|---|---|
| italic("E. coli") | no-bias | 0.2971968 | 0.0250915 |
| italic("E. coli") | pAT | 0.8785546 | 0.0000245 |
| italic("E. coli") | pGC | -0.7948746 | 0.0004635 |
| italic("S. pneumoniae") | no-bias | -0.0734709 | 0.7217002 |
| italic("S. pneumoniae") | pAT | 1.3010456 | 0.0000003 |
| italic("S. pneumoniae") | pGC | -1.0164955 | 0.0000001 |

```
### logY ~ logRSCU
p.RSCU.OR1 <- ggplot(OR_codons2, aes(log_rscu, log.OR1, label = codon_pair)) +
  geom_point(size = 0.6) +
  geom_smooth(method = "glm", formula = y~x, se = F, color = "black", size = 1) +
  xlab(expression(paste("log(", italic(RSCU[p]/RSCU[u]), ")", sep = ""))) +
  ylab(expression(paste("log(", italic("Y"), ")", sep = ""))) +
  stat_cor(label.x.npc = "left", label.y.npc = "bottom",
           method = "pearson", size = 3.5) +
  facet_wrap(species~mutation_type, scales = "free",
```

```r
               labeller = function(x) {x[2]}) +
  theme_bw() +
  theme(text = element_text(family = "Times", size = 14),
        axis.text = element_text(family = "Times", size = 14),
        strip.text.x = element_text(family = "Times", size = 14),
        strip.background.x = element_rect(fill = "gray90", linetype = "blank"))
p.RSCU.OR1
```

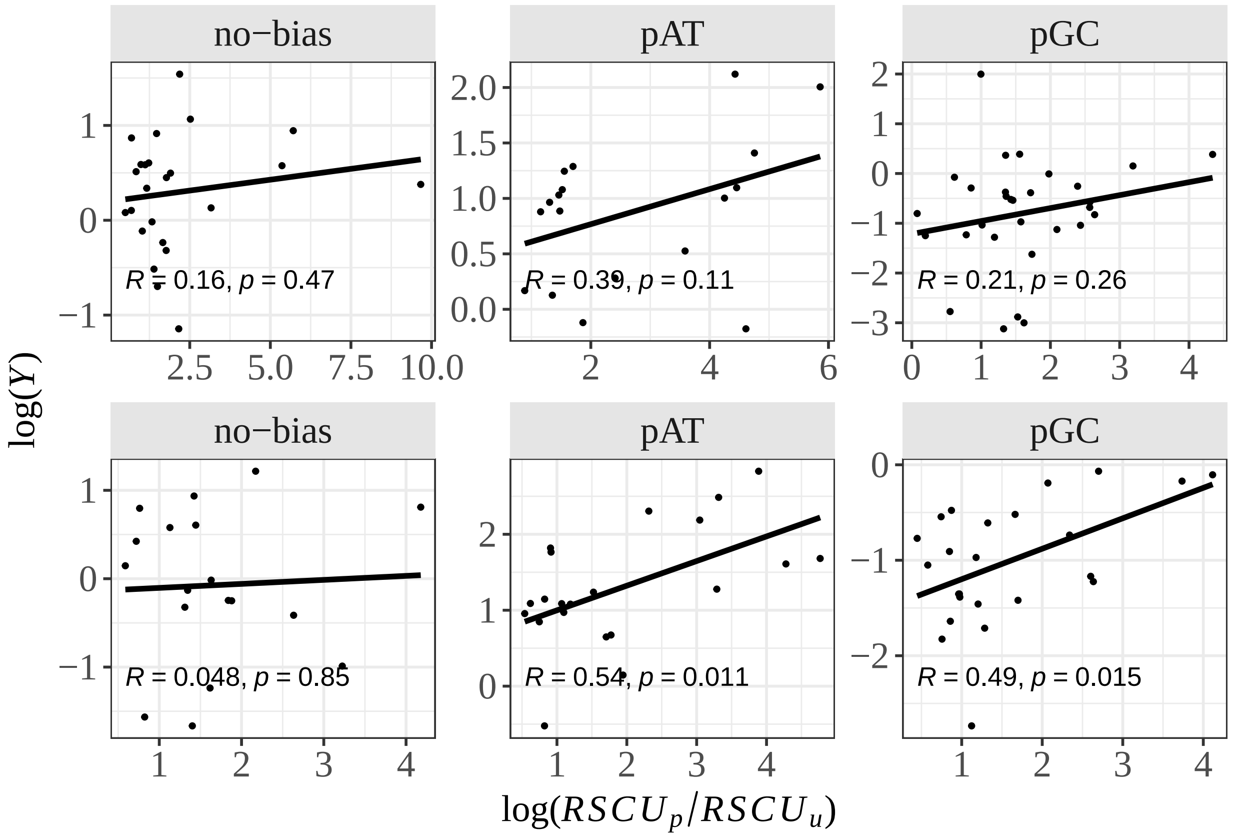


```r
# ANCOVA
lm_all <- lm(data = OR_codons2, formula = log.OR1~log_rscu+mutation_type+species)
kable(x = anova(lm_all))
```

| | Df | Sum Sq | Mean Sq | F value | Pr(>F) |
|---|---|---|---|---|---|
| log_rscu | 1 | 17.5754783 | 17.5754783 | 27.1738881 | 0.0000007 |
| mutation_type | 2 | 79.1359397 | 39.5679698 | 61.1770314 | 0.0000000 |
| species | 1 | 0.0187884 | 0.0187884 | 0.0290492 | 0.8649306 |
| Residuals | 130 | 84.0811651 | 0.6467782 | NA | NA |

```r
# linear model for each species:
ecoli_df <- subset(OR_codons2, species == 'italic("E. coli")')
spneu_df <- subset(OR_codons2, species == 'italic("S. pneumoniae")')
lm_ecoli <- lm(data = ecoli_df, formula = log.OR1~log_rscu+mutation_type)
lm_spneu <- lm(data = spneu_df, formula = log.OR1~log_rscu+mutation_type)
kable(x = anova(lm_ecoli))
```

|  | Df | Sum Sq | Mean Sq | F value | Pr(>F) |
|---|---|---|---|---|---|
| log_rscu | 1 | 8.512584 | 8.5125836 | 11.78387 | 0.0010212 |
| mutation_type | 2 | 28.311027 | 14.1555137 | 19.59532 | 0.0000002 |
| Residuals | 68 | 49.122703 | 0.7223927 | NA | NA |

```
kable(x = anova(lm_spneu))
```

|  | Df | Sum Sq | Mean Sq | F value | Pr(>F) |
|---|---|---|---|---|---|
| log_rscu | 1 | 11.14324 | 11.1432388 | 22.38083 | 1.44e-05 |
| mutation_type | 2 | 54.29780 | 27.1488995 | 54.52767 | 0.00e+00 |
| Residuals | 59 | 29.37564 | 0.4978921 | NA | NA |

```
# both log(RSCUp/RSCUu) and mutation type are significant
```

Assessing the impact of mutation bias and biased gene conversion through differences in allele frequency across codon pairs.

```
# function to estimate the folded site frequency spectrum (minor allele frequency)
buildSFS.folded <- function(dat.bpp, n, ntot) {
  for (i in 1:nrow(dat.bpp)) {
    size <- ntot - dat.bpp[i, "MissingDataFrequency"]
    if (dat.bpp[i, "NbAlleles"] == 1) {
      if (size < n) {
        # Discard position
        dat.bpp[i, "Freq"] <- NA
      } else {
        dat.bpp[i, "Freq"] <- 0
      }
    } else if (dat.bpp[i, "NbAlleles"] > 2) {
      dat.bpp[i, "Freq"] <- NA
    } else {
      if (size < n) {
        # Discard position
        dat.bpp[i, "Freq"] <- NA
      } else if (size == n) {
        # Get SFS
        dat.bpp[i, "Freq"] <- dat.bpp[i, "MinorAlleleFrequency"]
      } else {
        # Performs projection
        # We sample alleles
        x <- c(rep(1, dat.bpp[i, "MinorAlleleFrequency"]),
               rep(0, dat.bpp[i, "MajorAlleleFrequency"]))
        y <- sample(x, size = n, replace = FALSE)
        dat.bpp[i, "Freq"] <- sum(y)
      }
    }
  }
  return(dat.bpp)
}

# separating the species to estimate the folded site frequency spectrum for each species
ecoli_codons <- subset(codons_poly, species == "ecoli")
spneu_codons <- subset(codons_poly, species == "spneumoniae")
```

```r
# downsampling sites to account for missing data frequency
# (sites with a lot of missig data will not be sampled)
ecoli_codons_maf <- ddply(ecoli_codons, c("coID", "Site"), buildSFS.folded, 81, 82)
spneu_codons_maf <- ddply(spneu_codons, c("coID", "Site"), buildSFS.folded, 15, 16)


# estimating the mean allele frequency
ecoli_codons_maf$maf <- ecoli_codons_maf$Freq/81
spneu_codons_maf$maf <- spneu_codons_maf$Freq/15

# combining the two tables:
codons_maf <- rbind(ecoli_codons_maf, spneu_codons_maf)

# difference in allele frequecy between u>p and p>u
dif_maf <- ddply(codons_maf, c("species", "aa3_pair", "codon_pair"), function(x) {
  maf_p <- mean(x$maf[x$minor_codon == "p"], na.rm = T)
  maf_np <- mean(x$maf[x$minor_codon == "np"], na.rm = T)
  data.frame(maf_p, maf_np)
})

dif_maf$dif_maf <- dif_maf$maf_p - dif_maf$maf_np
dif_maf$log.maf <- log(dif_maf$maf_np/dif_maf$maf_p)

# getting OR table to add dif_maf information:
OR_codons2 <- read.table(file = "~/Dropbox/CUB_supplementary_data/tables/OR_tbl2_codons.csv",
                         sep = "\t", header = T)

# combing the two tables:
OR_codons3 <- merge(OR_codons2, dif_maf, by = c("species", "aa3_pair", "codon_pair"))
# saving the table with all information at the codon level:
write.table(OR_codons3, file = "~/Dropbox/CUB_supplementary_data/tables/OR_tbl3_codons.csv",
            sep = "\t", col.names = T, row.names = F, quote = F)



### RESULTS


OR_codons3$species <- factor(OR_codons3$species, levels = c("ecoli", "spneumoniae"))
levels(OR_codons3$species) <- c(expression(italic("E. coli")),
                                expression(italic("S. pneumoniae")))

# dif_maf distribution across mutation types:
dif_maf_dist <- ggplot(OR_codons3, aes(x=dif_maf)) +
  geom_histogram(aes(y=..density..),
                 binwidth=.02,
                 colour="black", fill="white") +
  xlab("Differences in allele frequency") +
  ylab("Frequency") +
  geom_vline(data = ddply(OR_codons3, c("species", "mutation_type"),
                          summarise, avg = mean(dif_maf)),
             aes(xintercept=avg),linetype="dashed",
             color = "azure4", size=0.5) +
```

```
  facet_grid(species~mutation_type, scales = "free", labeller = label_parsed) +
  geom_text(data = ddply(OR_codons3, c("species", "mutation_type"),
                         summarise, avg = mean(dif_maf, na.rm = T)),
            aes(x = -0.2, y = 10, label = paste("mean = ", round(avg, digits = 3), sep = "")),
            hjust = 0,family = "Times", size = 4, color = "azure4") +
  theme_bw() +
  theme(text = element_text(family = "Times", size = 14),
        axis.text = element_text(family = "Times", size = 14),
        strip.text.x = element_text(family = "Times", face = "bold", size = 14),
        strip.text.y = element_text(family = "Times", face = "bold", size = 14),
        strip.background.x = element_rect(fill = "gray90", linetype = "blank"),
        strip.background.y = element_rect(fill = "gray90", linetype = "blank"))
dif_maf_dist
```

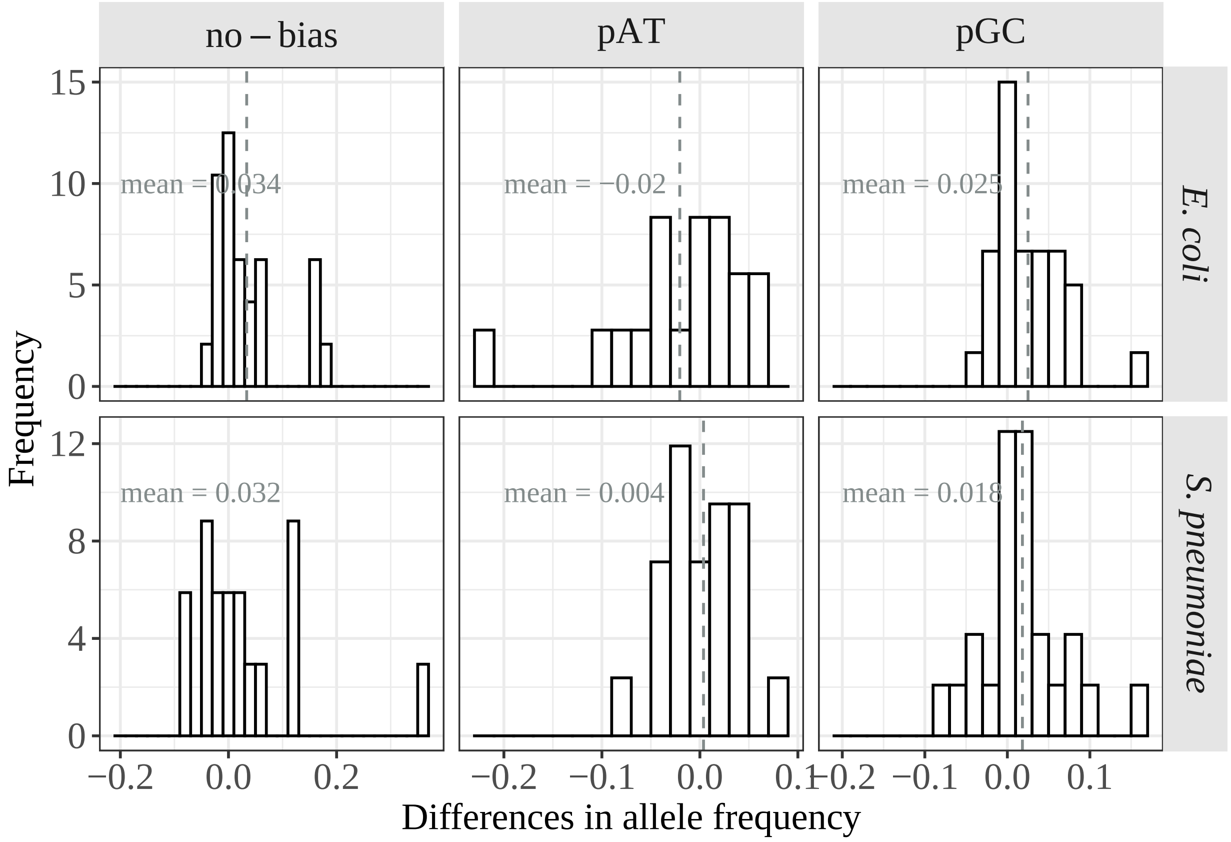


```
# checking if the mean is significantly different from 0:
P_maf <- ddply(OR_codons3, c("species", "mutation_type"), function(x) {
  t_test <- t.test(x$dif_maf)
  mean <- t_test$estimate
  p_value <- t_test$p.value
  data.frame(mean, p_value)
})
kable(P_maf)
```

| species | mutation_type | mean | p_value |
|---|---|---|---|
| italic("E. coli") | no-bias | 0.0337009 | 0.0170242 |
| italic("E. coli") | pAT | -0.0204951 | 0.2275399 |
| italic("E. coli") | pGC | 0.0250457 | 0.0024407 |
| italic("S. pneumoniae") | no-bias | 0.0319670 | 0.2357464 |
| italic("S. pneumoniae") | pAT | 0.0037140 | 0.6568853 |
| italic("S. pneumoniae") | pGC | 0.0182729 | 0.0957965 |

```
# ANOVA:
maf_lm <- lm(data = OR_codons3, formula = dif_maf~mutation_type+species)
kable(x = anova(maf_lm))
```

| | Df | Sum Sq | Mean Sq | F value | Pr(>F) |
|---|---|---|---|---|---|
| mutation_type | 2 | 0.0351686 | 0.0175843 | 4.528287 | 0.0125565 |
| species | 1 | 0.0004898 | 0.0004898 | 0.126121 | 0.7230638 |
| Residuals | 130 | 0.5048175 | 0.0038832 | NA | NA |

```
# separating species:
ecoli_maf <- subset(OR_codons3, species == 'italic("E. coli")')
lm_maf_ecoli <- lm(data = ecoli_maf, formula = dif_maf~mutation_type)
kable(x = anova(lm_maf_ecoli))
```

| | Df | Sum Sq | Mean Sq | F value | Pr(>F) |
|---|---|---|---|---|---|
| mutation_type | 2 | 0.0339271 | 0.0169636 | 5.173014 | 0.0080663 |
| Residuals | 69 | 0.2262677 | 0.0032792 | NA | NA |

```
spneu_maf <- subset(OR_codons3, species == 'italic("S. pneumoniae")')
lm_maf_spneu <- lm(data = spneu_maf, formula = dif_maf~mutation_type)
kable(x = anova(lm_maf_spneu))
```

| | Df | Sum Sq | Mean Sq | F value | Pr(>F) |
|---|---|---|---|---|---|
| mutation_type | 2 | 0.0075534 | 0.0037767 | 0.8170517 | 0.4466672 |
| Residuals | 59 | 0.2727176 | 0.0046223 | NA | NA |

```
## only significant in E. coli

# dif_maf ~ log(RSCU)
plot_rscu_maf <- ggplot(OR_codons3, aes(log_rscu, dif_maf, label = codon_pair)) +
  geom_point(size = 0.6) +
  geom_smooth(method = "glm", formula = y~x, se = F, color = "black", size = 1) +
  ylab("Differences in allele frequency") +
  xlab(expression(paste("log(", italic(RSCU[p]/RSCU[u]), ")", sep = ""))) +
  stat_cor(label.x.npc = "left", label.y.npc = "top", method = "pearson", size = 3.5) +
  facet_wrap(species~mutation_type, scales = "free", labeller = function(x) {x[2]}) +
  theme_bw() +
  theme(text = element_text(family = "Times", size = 14),
        axis.text = element_text(family = "Times", size = 14),
        strip.text.x = element_text(family = "Times", size = 14),
        strip.text.y = element_text(family = "Times", face = "bold", size = 14),
        strip.background.x = element_rect(fill = "gray90", linetype = "blank"),
        strip.background.y = element_rect(fill = "gray90", linetype = "blank"))
plot_rscu_maf
```

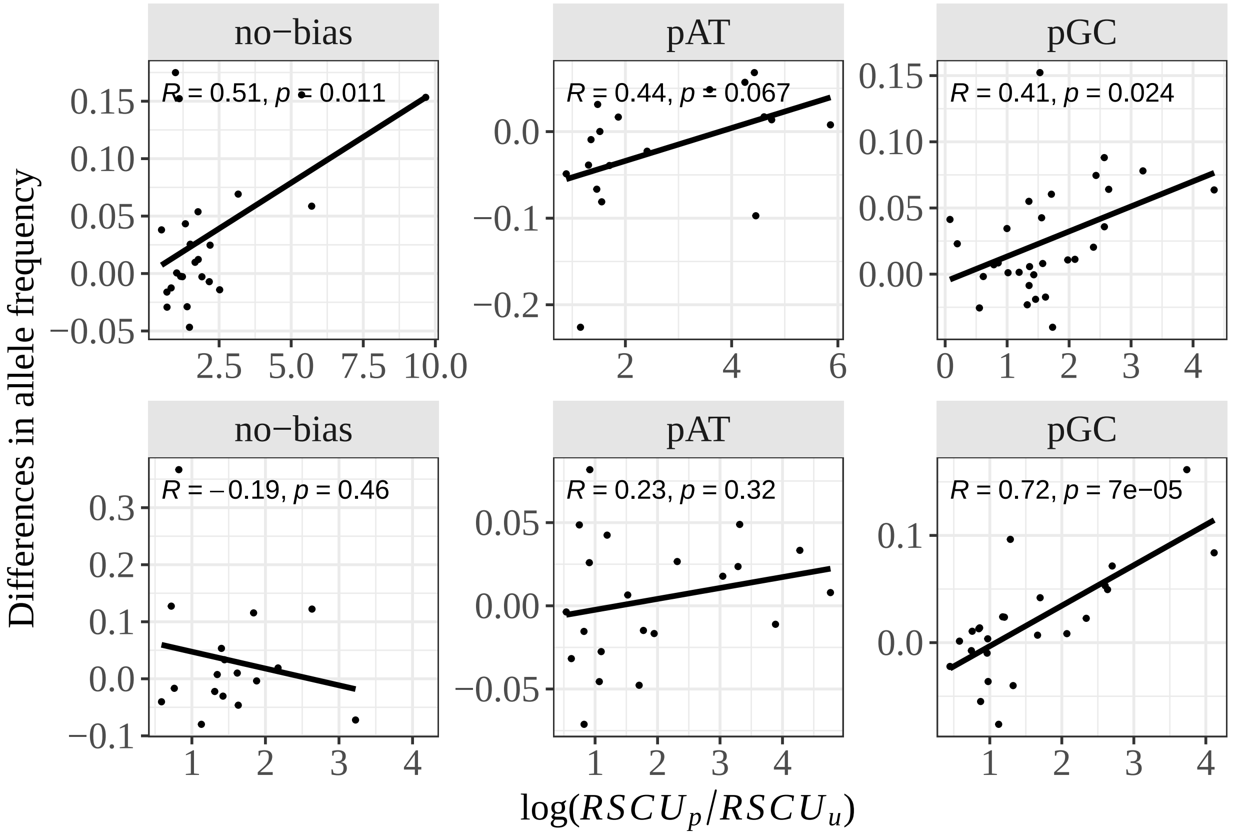


The following script contains the analysis of Z while considering the type of mutation.

```
# table with codon substitutions and RSCU values from highly expressed genes
codon_subs_high <- fread(file = "~/Dropbox/CUB_supplementary_data/tables/codon_subs_rscu_high.csv",
                         sep = "\t", header = T)

# including the mutation in each aa pair:
codon_subs_high$codon2 <- as.character(codon_subs_high$codon2)
codon_subs_high$codon1 <- as.character(codon_subs_high$codon1)

# function that adds the mutation type in the table:
nt_change.fun <- function(tbl) {
  for (i in 1:nrow(tbl)) {
    aa1 <- strsplit(tbl[i, "codon2"], split = "")
    aa2 <- strsplit(tbl[i, "codon1"], split = "")
    comp.aa <- stri_compare(aa1[[1]], aa2[[1]])
    j <- which(!comp.aa == 0)
    tbl[i, "mutation"] <- paste(sort(c(aa1[[1]][j], aa2[[1]][j]))[1],
                                sort(c(aa1[[1]][j], aa2[[1]][j]))[2], sep = "")
  }
  return(tbl)
}

codon_subs_high$pos <- seq(1:nrow(codon_subs_high))

# applying the function on the dataset:
```

```
df_subs_codons <- ddply(codon_subs_high, "pos", nt_change.fun)

# considering only non-synoynous substitutions seperated by 1 mutational step
df_subs_ns_codons <- subset(df_subs_codons, nt.dif == 1 & !(aa1 == aa2))

# adding codon pair to the table:
temp.codon <- cbind(t(apply(df_subs_ns_codons[,c(1,2)], 1, sort)))
df_subs_ns_codons$codon_pair <- paste(temp.codon[,1], temp.codon[,2], sep = "-")

# getting the p and u codons from the OR table (polymorphism table):
OR_codons3 <- read.table(file = "~/Dropbox/CUB_supplementary_data/tables/OR_tbl3_codons.csv",
                         sep = "\t", header = T)

# taking only the columns we need:
codon_type <- unique(subset(OR_codons3,
                            select = c("codon_pair", "p_codon", "u_codon", "p_allele",
                                       "u_allele", "mutation_type", "species")))

# merging with the substitutionstable with CAI information
df2_subs_ns_codons <- merge(df_subs_ns_codons, codon_type, by = c("codon_pair", "species"))

# estimating the weighted averages:
df2_subs_ns_codons$weight1 <- df2_subs_ns_codons$counts1*(log((df2_subs_ns_codons$rscu2 + 1e-04)/
                                                                (df2_subs_ns_codons$rscu1 + 1e-04)))

# estimating Z for each amino acid in each mutation type:
mutation_type_Z <- ddply(df2_subs_ns_codons, c("species", "mutation_type", "aa_pair"),
                         function(x) {
  Z = sum(x$weight1, na.rm = TRUE)/sum(unique(x$counts1), na.rm = TRUE)
  data.frame(Z)
})

# adding dxy information:
dxy_df <- read.table(file = "~/Dropbox/CUB_supplementary_data/tables/dxy_df.csv",
                     sep = "\t", header = T)
mutation_type_Z <- merge(mutation_type_Z, dxy_df, by = c("species", "aa_pair"))

# Z distribution across mutation types:
mutation_type_Z$species <- factor(mutation_type_Z$species,
                                  levels = c("ecoli", "spneumoniae"))
levels(mutation_type_Z$species) <- c(expression(italic("E. coli")),
                                     expression(italic("S. pneumoniae")))

p.dist.Z_mutation <- ggplot(mutation_type_Z, aes(x=Z)) +
  geom_histogram(aes(y=..density..),
                 binwidth=.3,
                 colour="black", fill="white") +
  xlab(expression(italic(Z))) +
  ylab("Frequency") +
  geom_vline(data = ddply(mutation_type_Z, c("species", "mutation_type"),
                          summarise, avg = mean(Z)),
             aes(xintercept=avg),linetype="dashed",
             color = "azure4", size=0.5) +
```

```
  facet_grid(species~mutation_type, scales = "free", labeller = label_parsed) +
  geom_text(data = ddply(mutation_type_Z, c("species", "mutation_type"),
                        summarise, avg = mean(Z)),
           aes(x = -7, y = 0.9, label = paste("mean(Z) = ",
                                             round(avg, digits = 3), sep = "")),
           hjust = 0,family = "Times", size = 4, color = "azure4") +
  theme_bw() +
  theme(text = element_text(family = "Times", size = 14),
        axis.text = element_text(family = "Times", size = 14),
        strip.text.x = element_text(family = "Times", size = 14),
        strip.text.y = element_text(family = "Times", face = "bold", size = 14),
        strip.background.x = element_rect(fill = "gray90", linetype = "blank"),
        strip.background.y = element_rect(fill = "gray90", linetype = "blank"))
p.dist.Z_mutation
```

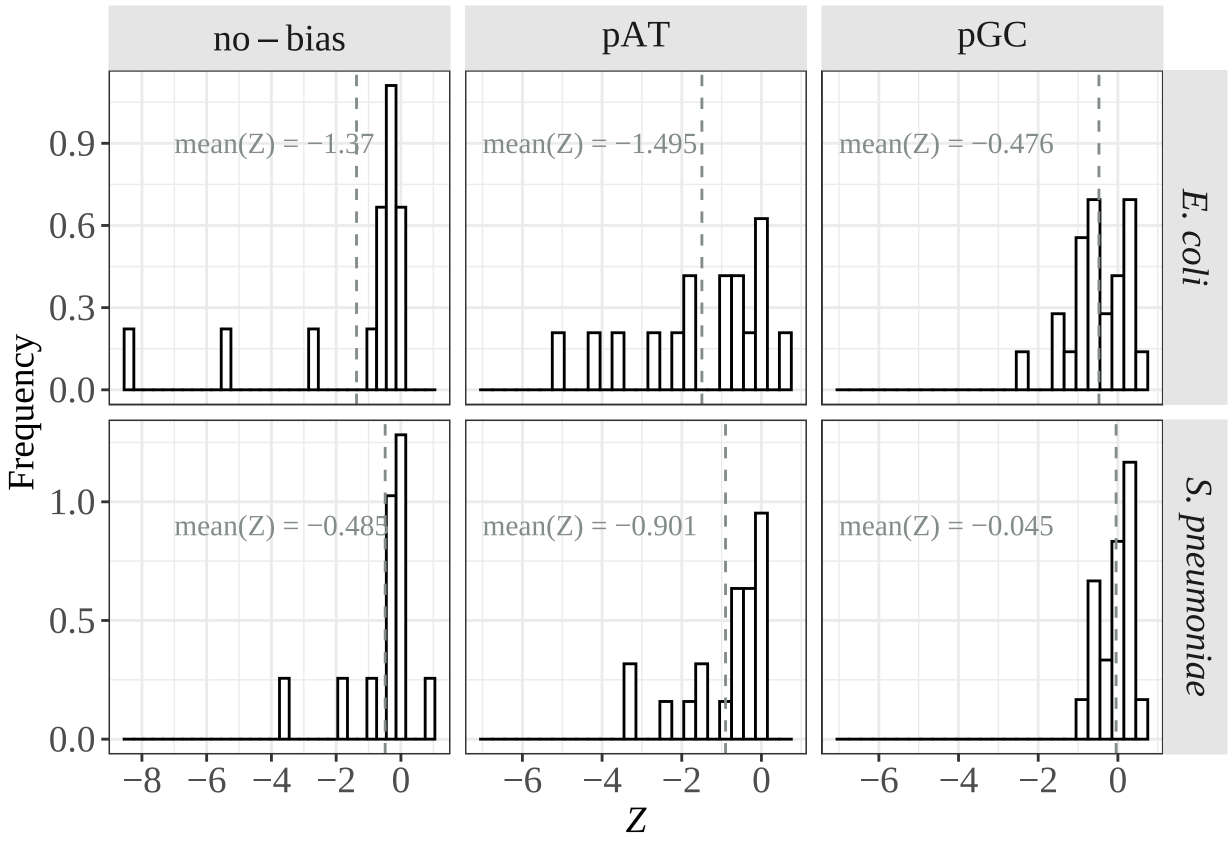


```
# Z ~ dxy + mutation type
p.Z.dxy_mutation <- ggplot(mutation_type_Z,
                           aes(Z, dxy, label = aa_pair)) +
  geom_point(size = 0.6) +
  geom_smooth(method = "glm", formula = y~x, se = F, color = "black", size = 1) +
  xlab(expression(italic(Z))) +
  ylab(expression(italic(d[ij]))) +
  stat_cor(label.x.npc = "left", label.y.npc = "top", method = "pearson", size = 3.5) +
  facet_wrap(species~mutation_type, scales = "free", labeller = function(x) {x[2]}) +
  theme_bw() +
```

```r
  theme(text = element_text(family = "Times", size = 14),
        strip.text.x = element_text(family = "Times", size = 14),
        strip.text.y = element_text(family = "Times", face = "bold", size = 14),
        strip.background.x = element_rect(fill = "gray90", linetype = "blank"),
        strip.background.y = element_rect(fill = "gray90", linetype = "blank"))
p.Z.dxy_mutation
```

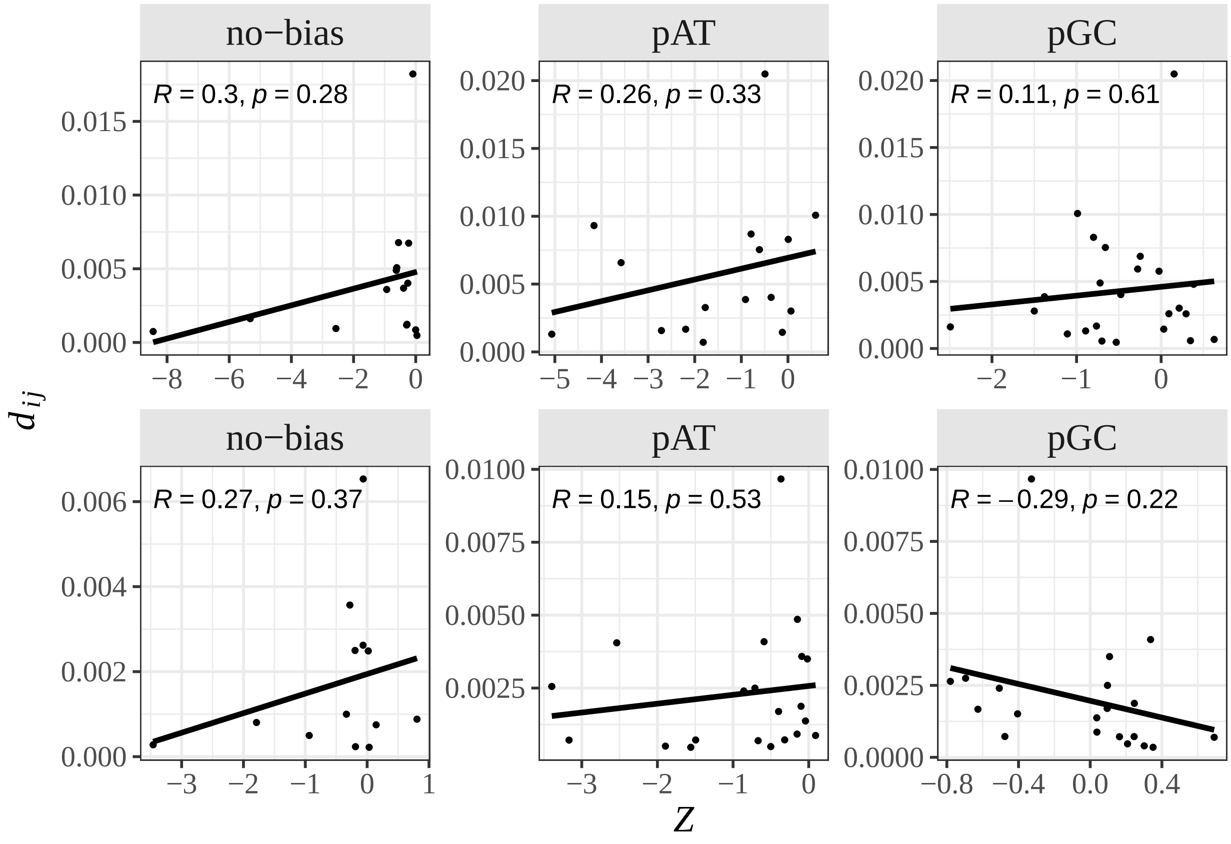


```r
# top: ecoli
# bottom: spneumoniae
```
